# The E3 Ubiquitin Ligase Nedd4L Acts as a Checkpoint Against Activation in Quiescent Muscle Stem Cells

**DOI:** 10.1101/2023.05.10.540205

**Authors:** Darren M. Blackburn, Korin Sahinyan, Aldo Hernández-Corchado, Felicia Lazure, Vincent Richard, Laura Raco, René P. Zahedi, Christoph H. Borchers, Christoph Lepper, Hiroshi Kawabe, Arezu Jahani-Asl, Hamed S. Najafabadi, Vahab D. Soleimani

**Affiliations:** Department of Human Genetics, McGill University, 3640 rue University, Montréal, QC, H3A 0C7, Canada; Lady Davis Institute for Medical Research, Jewish General Hospital, 3755 Chemin de la Côte-Sainte-Catherine, Montréal QC, H3T 1E2, Canada; Segal Cancer Proteomics Centre, Lady Davis Institute, Jewish General Hospital, McGill University, Montréal, Quebec H3T 1E2, Canada; Manitoba Centre for Proteomics and Systems Biology, Winnipeg, MB R3E 3P4, Canada; Department of Internal Medicine, University of Manitoba, Winnipeg, MB R3E 3P4, Canada; Department of Biochemistry and Medical Genetics, University of Manitoba, Winnipeg, MB R3E 0J9, Canada; Gerald Bronfman Department of Oncology, Lady Davis Institute for Medical Research, Jewish General Hospital, Montréal, QC H3T 1E2, Canada; Division of Experimental Medicine, McGill University, Montréal, QC H4A 3J1, Canada; Department of Pathology, McGill University, Montréal, QC H3A 2B4, Canada; Department of Physiology & Cell Biology, College of Medicine, The Ohio State University, Columbus, OH, USA; Department of Molecular Neurobiology, Max Planck Institute of Experimental Medicine, 37075, Göttingen, Germany; Department of Cellular and Molecular Medicine and University of Ottawa Brain and Mind Research Institute, University of Ottawa, Ottawa, ON, Canada

**Keywords:** Stem cell quiescence, Skeletal muscle stem cells, Transcriptomics, Proteomics, Nedd4L, Ubiquitin proteasome system

## Abstract

Adult stem cells play a critical role in tissue repair and maintenance. In tissues with slow turnover, including skeletal muscle, these cells are maintained in a mitotically quiescent state yet remain poised to re-enter the cell cycle to replenish themselves and regenerate the tissue. Using a multiomics approach we identify the PAX7/NEDD4L axis as a checkpoint against muscle stem cell activation in homeostatic skeletal muscle. Our findings demonstrate that PAX7 transcriptionally activates the E3 ubiquitin ligase Nedd4L and that the conditional genetic deletion of Nedd4L impairs muscle stem cell quiescence, with an upregulation of cell cycle and myogenic differentiation genes. Loss of Nedd4L in muscle stem cells results in the expression of DCX which is only expressed during their *in vivo* activation. Together, this data establishes that the ubiquitin proteasome system, mediated by Nedd4L, is a key regulator of the muscle stem cell quiescent state in non-injured skeletal muscle.

**Highlights:**

- General inhibition of the ubiquitin proteasome system with MG132 results in muscle stem cells (MuSCs) breaking quiescence.
- The E3 ubiquitin ligase Nedd4L is a transcriptional target of Pax7.
- The Pax7/Nedd4l axis restricts MuSC activation in homeostatic skeletal muscle.
- Genetic deletion of Nedd4L induces MuSCs’ transition towards activation.

## Introduction

Tissue specific stem cells are essential for the maintenance and regeneration of the tissue in which they reside. Postnatally, adult skeletal muscle is maintained by a rare population of muscle stem cells (MuSCs), also known as satellite cells ^1, 2^. In the absence of trauma or injury, these cells typically exist in a quiescent state and are marked by the expression of the transcription factor Pax7 ^3, 4^. However, they can quickly respond in the event of an injury to the muscle and activate to re-enter the cell cycle. Once activated, MuSCs will proliferate, differentiate and self-renew, thus repairing the damaged muscle and restoring tissue homeostasis ^2^.

Maintenance of quiescence is crucial to prevent the exhaustion of the adult stem cell pool to precocious activation and differentiation. Previous studies have demonstrated that stem cell quiescence is a complex and dynamic state which is actively maintained ^5^. For example, the state of G_alert_ was first described in MuSCs ^6^ in which the stem cells are no longer in deep quiescence but have not yet fully transitioned into activation and the cell cycle. Differences in the depth of quiescence were also identified in other adult stem cells, including neuronal stem cells (NSC) and hematopoietic stem cells (HSC) ^7–9^. Characteristics of G_alert_ include larger cell size, higher mitochondria content, increased translation and a more rapid cell division at the time of activation ^6^. Multiple signalling pathways have been identified to facilitate the transition of MuSCs into G_alert_, including the activation of mTORC1 ^6^, the genetic deletion of the mitochondrial protein OPA1 ^10^ and GLI3 ^11^ or an injury to a distant tissue ^12^. Together, these studies suggest that transition of quiescent adult stem cells into cell cycle constitutes a continuum with potentially numerous checkpoints along the way.

In adult stem cells, several mechanisms have been identified which poise the cells for activation while simultaneously maintaining quiescence^13^. Of note, in MuSCs it is known that there is a large amount of bivalent chromatin, possessing both permissive and repressive marks, indicating that these genes can be quickly transcribed in the case of activation^14^. MuSCs also have low mRNA content due primarily to transcriptional repression from the lack of RNA polymerase II phosphorylation^15, 16^. Translational control of the produced mRNA can play a role in the poised state of MuSCs. It has been previously reported that both Myf5 and MyoD1 transcripts can be sequestered in quiescent MuSCs, preventing their translation^17, 18^. mRNA degradation by miRNAs is also a well described mechanism for the maintenance of quiescence, with these miRNAs quickly downregulated in the case of activation^19, 20^.

The ubiquitin proteasome system (UPS) is responsible for the targeted degradation of proteins. Proteins that are destined for degradation will be K48 polyubiquitinated. This process is comprised of a chain of enzymatic reactions executed by the E1 ubiquitin activating enzyme, an E2 ubiquitin transporter, and an E3 ubiquitin ligase^21^. The role of the UPS in the maintenance of MuSC quiescence is not well understood, however, the disruption of the UPS via the genetic deletion of *Rpt3*, a component of the proteasome, has been shown to cause the MuSCs to become apoptotic^22^. In HSCs undergoing activation, select E3 ubiquitin ligases have been shown to decrease in expression ^23–25^, which could suggest a role for these enzymes in the regulation of adult stem cell quiescence. A recent study has shown that the Skp1-Cul1-F-box protein ubiquitin ligase complex, maintains MuSC quiescence ^26^.

Key members of the UPS are the E3 ubiquitin ligases. These enzymes are responsible for tagging the proteins with ubiquitin, targeting them to the proteasome for degradation. Neural Precursor Cell Expressed Developmentally Downregulated 4-Like (Nedd4L) is an E3 ubiquitin ligase that is part of the HECT family^27^. Nedd4l is known to target a number of membrane proteins, such as the amiloride-sensitive epithelial Na+ channel (ENaC), TGFBR1, and voltage gated sodium channels (Na_v_s)^28–32^. It is also known to act as a tumor suppressor in human cancers, with a decrease in its expression correlating with worse prognosis^27, 33–36^.

The role of Nedd4l in the context of MuSC quiescence has not been investigated. In the present study we report a central role for Nedd4l in maintaining MuSC quiescence downstream of PAX7. The PAX7/NEDD4L axis acts as a checkpoint between MuSC quiescence and activation. Importantly, genetic deletion of Nedd4l results in the MuSCs to exit quiescence and entry into G_alert_.

## Results

### Pax7 transcriptionally regulates the expression of the E3 ubiquitin ligase Nedd4L

To confirm that the UPS is essential for maintaining MuSC quiescence, we injected tibialis anterior (TA) muscles of mice with MG132, a general inhibitor of the proteasome system, for 3 days (Figure 1A). Analysis of the TA cross sections showed that the MuSCs in the treated TAs were more prone to exit quiescence as shown by the expression of the proliferation marker Ki67 (Figure 1A-D). RNA-Seq analysis of freshly sorted MuSCs and primary myoblasts showed that 252 E3 ubiquitin ligases are expressed at a threshold of at least 100 base counts. An unbiased analysis of the E3 ubiquitin ligases whose expression change during the transition from quiescent muscle stem cells to cultured primary myoblasts, revealed that 120 E3 Ligases exhibit a significant change in expression (Supplemental Figure 1). We chose to focus on Nedd4L as it is highly expressed in quiescent MuSCs and highly conserved in mammals. Nedd4L is part of the Nedd4 family of E3 ubiquitin ligases. While Nedd4 is very highly expressed in muscle tissue, its expression is relatively static throughout the myogenic pathway and is not differentially expressed between quiescent muscle stem cells and myoblasts (Supplemental Figure 1B). Through RNA-Seq, we see that Nedd4L is most highly expressed in quiescent MuSCs, with a drop in expression in the myoblast stage and an increase in expression in the terminally differentiated myofibers (Figure 1E). Additionally, the expression profile of Nedd4L is highly dynamic in myofiber-associated MuSCs during *in vitro* culture (Figure 1F-G).

**Figure 1:**
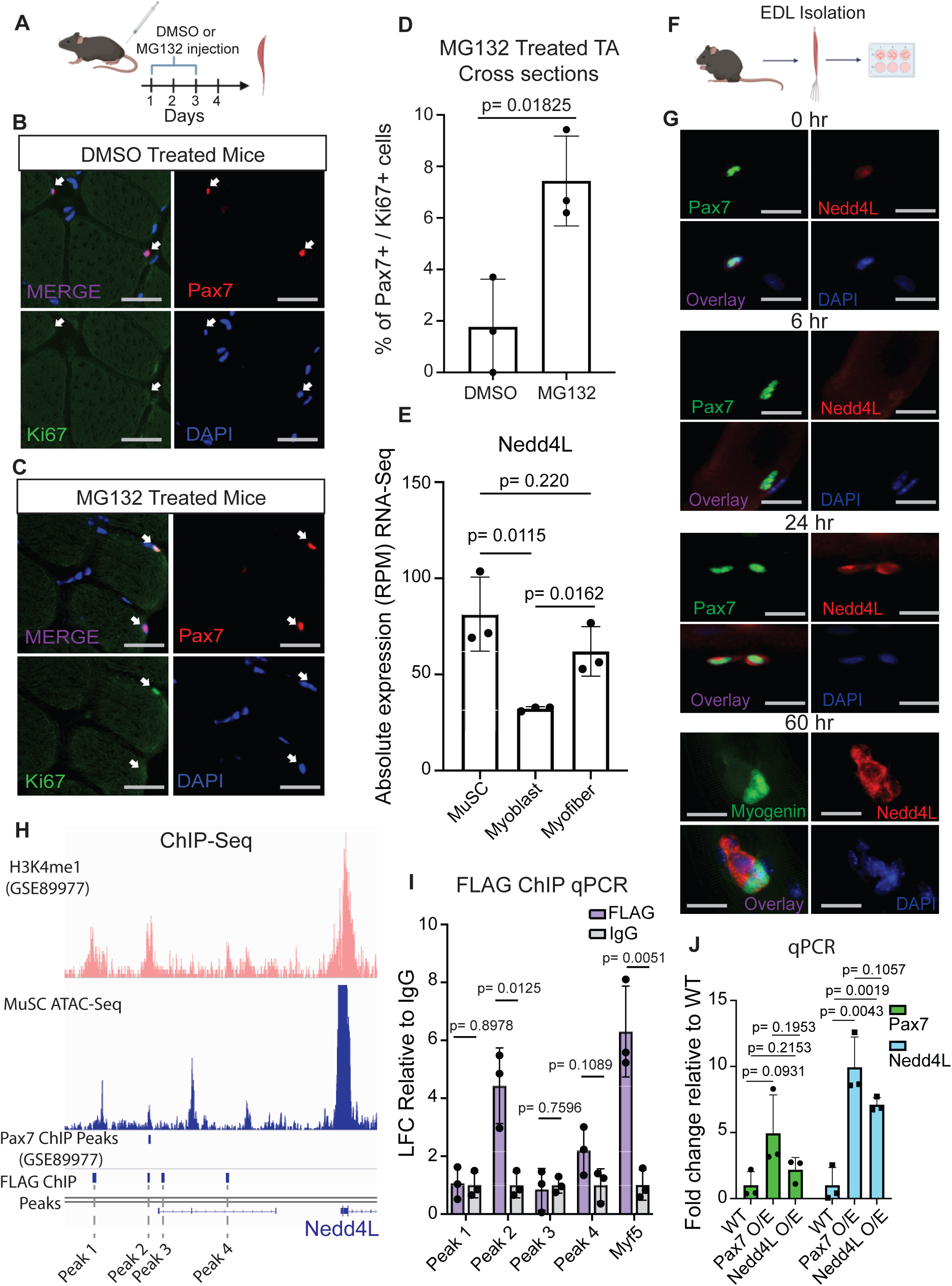
Pax7 transcriptionally regulates the expression of the E3 ubiquitin ligase Nedd4L. **A.** Diagram showing the TA of WT mice were intramuscularly injected with either DMSO or MG132 for 3 days, generated with Biorender.com. **B.** Representative immunofluorescent staining for Pax7 and Ki67 of cross sections from TA muscle treated with DMSO control. **C.** Representative immunofluorescent staining for Pax7 and Ki67 of cross sections from TA muscle treated with MG132. **D.** Bar graph representing the percentage of double positive Pax7^+^ and Ki67^+^ MuSCs. **E.** Bar graph of the absolute expression from RNA-Seq of Nedd4L from freshly isolated MuSCs, myoblasts three days post isolation, and single EDL myofibers. Single myofiber data is retrieved from the GEO GSE138591^67^. **F.** Diagram of EDL isolation from mice, generated with Biorender.com. **G.** Immunofluorescent staining of EDL myofibers for Pax7 and Nedd4L at 0, 6, and 24 hours post isolation, and staining for Myogenin and Nedd4L at 60 hours post isolation. **H.** IGV tracks of the enhancer region upstream of the Nedd4L promotor. The H3K4me1 tracks (GSM2394788) were retrieved from the GEO GSE89977^38^ and the Pax7 ChIP Peaks were called from the signal of GSM2394784 (+dox) by using GSM2394785 (-dox) as background ^38^. The Pax7-FLAG ChIP peaks were generated from primary myoblasts that were over expressing Pax7-FLAG. **I.** Bar graph of the FLAG ChIP qPCR of Pax7-FLAG overexpressing primary myoblasts, representing the LFC enrichment relative to the IgG control of the four nearest PAX7 peaks as determined by the FLAG ChIP-Seq with Myf5 as a positive control. **J.** qPCR of Pax7 and Nedd4L of WT, Pax7-FLAG overexpressing and Nedd4L overexpressing primary myoblasts. Data presented as mean ± SD.

Pax7 is a critical transcription factor for MuSC function and is highly expressed in quiescent MuSCs^3, 37^. Therefore, we analyzed independent ChIP-seq datasets for potential regulation of Nedd4L by Pax7 in muscle cells. ChIP Sequencing data from Lilja et al. indicate that there is a proximal enhancer region upstream from the Nedd4L promoter^38^ (Figure 1H). The study by Lilja et al. used a doxycycline system to express Pax7 in pluripotent stem cells and from their Pax7 ChIP-Seq we see that there is a Pax7 binding peaks at these enhancers^38^, which is confirmed with our own Pax7-FLAG ChIP-Seq in myoblasts (Figure 1H). These enhancers are also accessible in MuSCs as seen by our ATAC-Seq data on freshly sorted MuSCs (Figure 1H). Together these data suggests that Pax7 regulates Nedd4L expression. To further confirm this finding, we generated myoblasts stably over expressing Pax7-FLAG protein and performed a FLAG ChIP-qPCR. We observed that there is enrichment for PAX7 binding to theses enhancer elements (Figure 1I). Next, with real time qPCR of the RNA from the Pax7-FLAG overexpressing myoblasts we confirmed the upregulation of Nedd4L (Figure 1J). These data indicate that Pax7 positively regulates the expression of Nedd4L.

### Genetic Deletion of Nedd4L Results in the Loss of Quiescence-Associated Genes in MuSCs

To further elucidate the role of Nedd4L in MuSC function, we used a genetic mouse model where Nedd4L is specifically deleted in MuSCs using a Cre-loxP system. In this model, exon 15 of *NEDD4L*, which codes for part of the catalytic domain of the protein, is flanked by loxP sites and deleted in cells expressing Pax7 (Supplemental Figure 2A). The gene is still being transcribed, but without Exon 15 it is catalytically inactive (Supplemental Figure 2A-B).

We analyzed the transcriptome of freshly isolated MuSCs by Fluorescence Activated Cell Sorting (FACS) from WT and Nedd4L-cKO mice using SMART-Seq (Figure 2A). From this analysis we found that 426 genes were differentially expressed between WT and Nedd4L-cKO mice (Figure 2B). Principal Component Analysis (PCA) shows a clear separation of WT and Nedd4L-cKO MuSCs, and with Pearson correlation we note that Nedd4L-cKO cells are more similar to one another than their WT counterparts (Figure 2C-E). Interestingly, some of the most significantly downregulated genes, such as *NR1D1*, *CHRDL2* and *CALCR* are known markers of quiescence (Figure 2F-H). *NR1D1* is known to inhibit proliferation and myogenesis^39^, and is greatly reduced in expression in myoblasts compared to MuSCs (Figure 2F). *CALCR* is a receptor that has been determined to be important for the maintenance of MuSC quiescence^40, 41^. *CHRDL2* is known to be a marker of quiescence^42, 43^ and whose expression is completely lost in myoblasts (Figure 2G). The loss of *Chrdl2* transcript in the Nedd4L-cKO correlates with a reduction in its protein levels in freshly isolated EDL myofiber associated MuSCs of Nedd4L-cKO mice (Figure 2I). Further analysis of the transcriptomic data shows that of the top 100 most differentially expressed genes, the majority are upregulated (Supplemental Figure 2C). We questioned whether this was due to changes in RNA stability and found that only 10 genes have a change in their stability. Interestingly, 2 of the genes that had a decrease in their RNA stability are *NR1D1* and *PAX7* (Supplemental Figure 2D), which may explain the loss of *Nr1d1* expression in the Nedd4L-cKO MuSCs.

**Figure 2:**
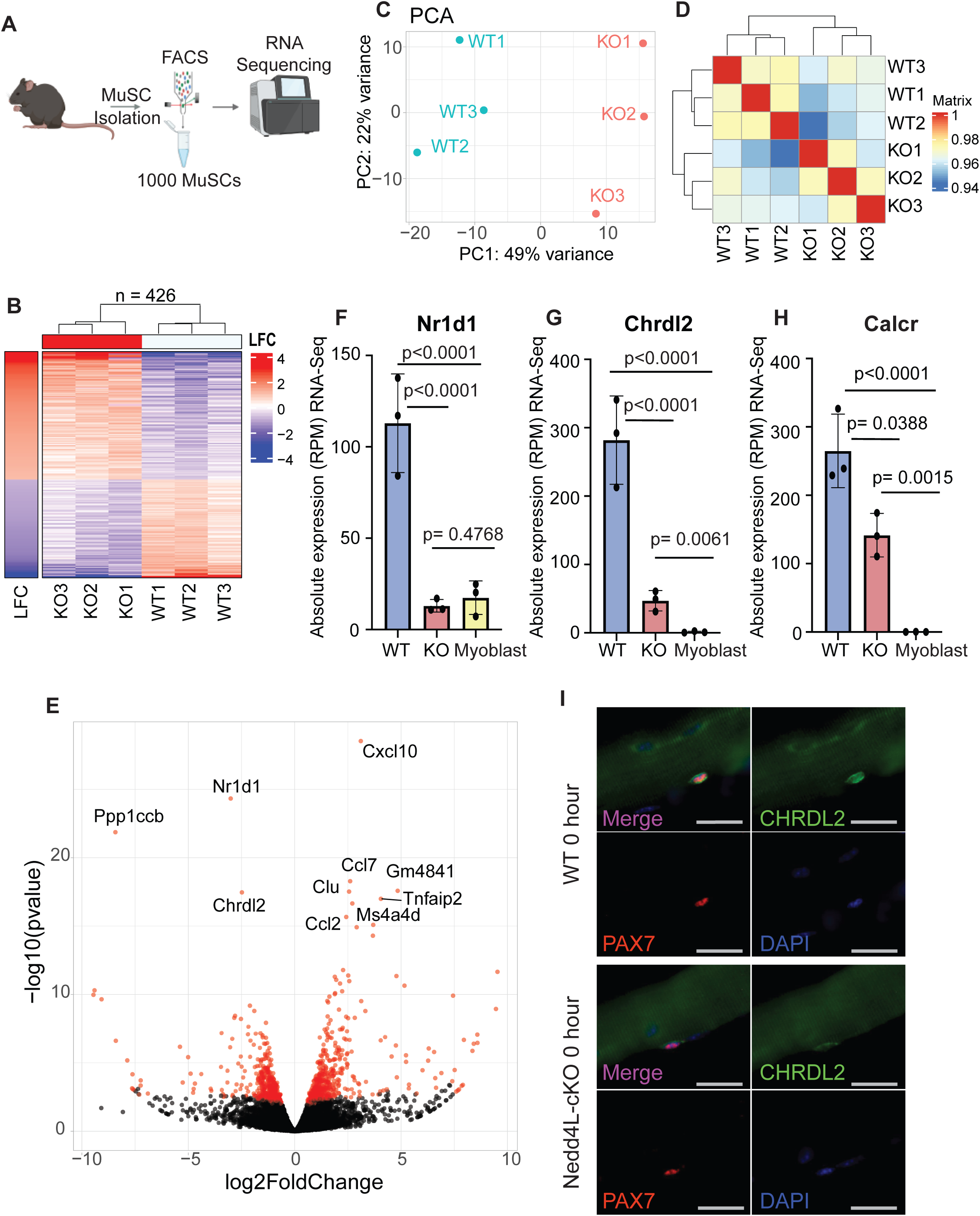
Genetic Deletion of Nedd4L Results in the Loss of Quiescence-Associated Genes in MuSCs. **A**. Schematic of the isolation of muscle stem cells and RNA-Seq, generated with Biorender.com. **B**. Heatmap of differentially expressed genes between WT and Nedd4L-cKO MuSCs. Threshold is adjusted p value ≤ 0.05, LFC ≥ 1, RPM ≥ 5. **C**. PCA plot of WT and Nedd4L-cKO MuSCs. **D**. Pearson correlation of the WT and Nedd4L-cKO MuSCs. **E**. Volcano plot of WT vs Nedd4L-cKO transcripts. **F**. Bar graph of the absolute expression (RPM) of *Nr1d1* in WT and Nedd4L-cKO MuSCs and in WT myoblasts. **G**. Bar graph of the absolute expression (RPM) of *Chrdl2* in WT and Nedd4L-cKO MuSCs and in WT myoblasts. **H**. Bar graph of the absolute expression (RPM) of *Calcr* in WT and Nedd4L-cKO. **I.** Immunofluorescence of Pax7 and Chrdl2 in MuSCs associated to freshly isolated EDL myofibers. Scale bar = 25 µm. Data presented as mean ± SD.

When we further analyse the pathways associated with the deregulated genes, we see that the majority of the dysregulated Reactome Pathways are upregulated (Figure 3A-B). Of the 53 pathways that are deregulated at a threshold of p-adj<0.1, 47 of them are upregulated compared to 6 that are downregulated (Figure 3A-B). Interestingly, many of the pathways that are upregulated are associated with MuSC activation, namely Translation, Respiratory Electron Transport, Cell Cycle Checkpoints and Muscle Contraction, among others (Figure 3 A,C).

**Figure 3:**
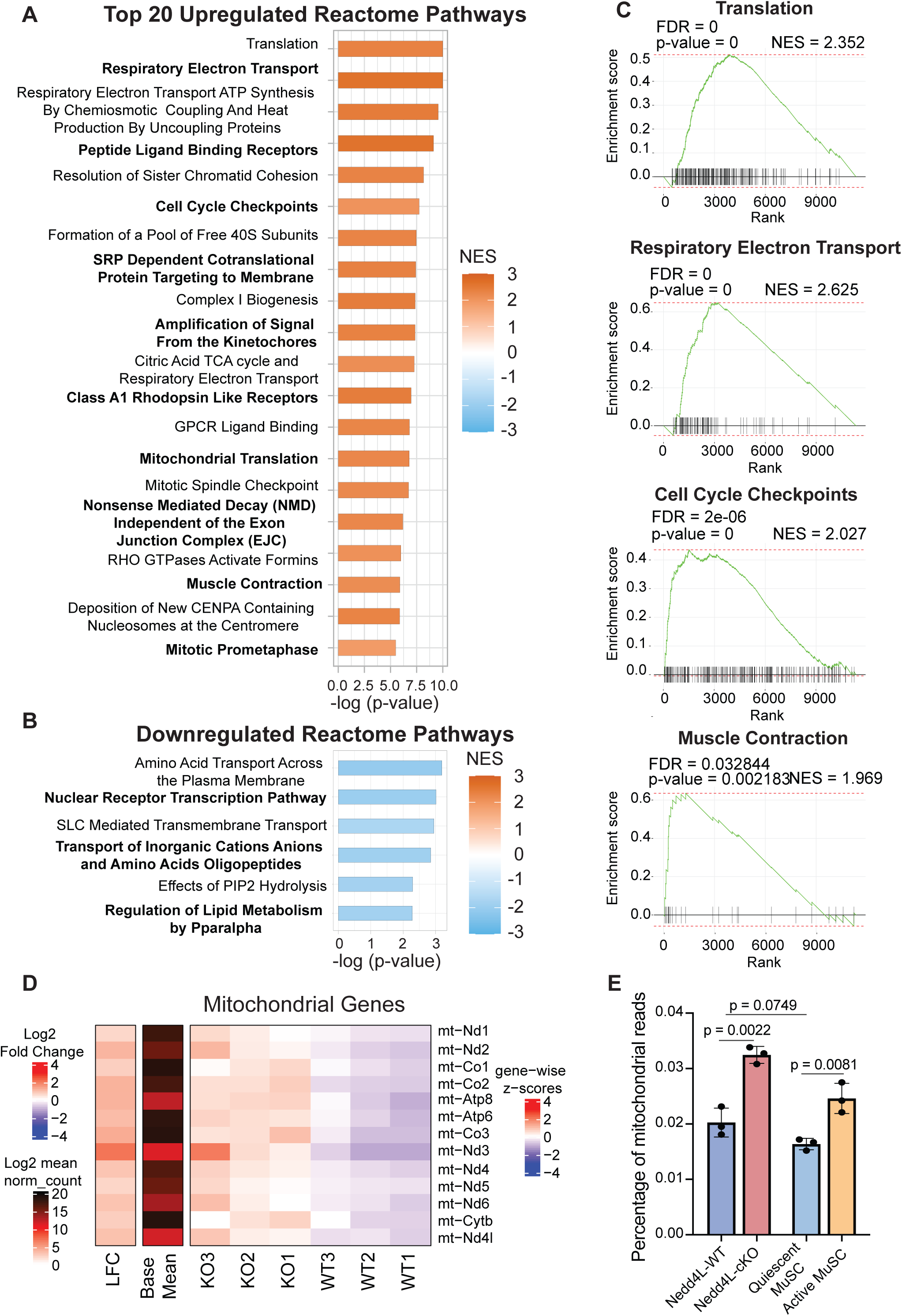
Genetic deletion of Nedd4L upregulates pathways associated with activation and differentiation. **A**. Top 20 upregulated Reactome pathways in Nedd4L-cKO MuSCs. **B**. All downregulated Reactome pathways in Nedd4L-cKO MuSCs. **C.** Selected enrichment plots for the Translation, Respiratory Electron Transport, Cell Cycle Checkpoints, and Muscle Contraction pathways. **D.** Heatmap of the expression of the mitochondrial genes in WT and Nedd4L-cKO MuSCs. **E.** Bar graph of the percentage of mitochondrial reads from the RNA-Seq of WT and Nedd4L-cKO MuSCs, with the quiescent and activated muscle stem cell RNA-Seq is from publicly available data retrieved from the GEO GSE144871^11^. Data presented as mean ± SD.

A similar trend is seen with the Hallmark Pathways where we see that the majority of the pathways are upregulated in the Nedd4L-cKO MuSCs (Supplemental Figure 3A). Likewise, many of the pathways identified are associated with MuSC activation and differentiation. Here we highlight the Myogenesis, G2M checkpoint, and Il6-Jak-Stat3 pathways (Supplemental Figure 3A-B). When we delve further into the genes that are affiliated with these pathways, we identify numerous genes that had previously been reported to be important for late myogenic differentiation, such as *MYMK, TNNT3,* and *Myl1* among others, are upregulated in the Nedd4L-cKO MuSCs (Supplemental Figure 3C). Additionally, a number of cell cycle related genes are upregulated in the Nedd4L-cKO condition, for example *CDK1, AURKA,* and *BIRC5* which are known to be downregulated in quiescent stem cells^44, 45^ (Supplemental Figure 3C). The Jak/Stat pathway has multiple roles in muscle, and its upregulation is associated with both proliferation and differentiation of muscle stem cells^46–50^. We see that *IL6* expression is upregulated in the Nedd4L-cKO MuSCs, along with other inflammatory cytokines (Supplemental Figure 3C). Interestingly, every mitochondrial gene in the Nedd4L-cKO MuSCs was significantly upregulated, correlating with the upregulation of the oxidative phosphorylation pathway (Figure 3D, Supplemental Figure 3A). A previous study has shown that there is a metabolic shift in MuSCs during activation, that MuSCs will increase their oxidative phosphorylation^51^. We also observe an increase in the percentage of mitochondrial reads from the RNA-seq of WT and Nedd4L-cKO MuSCs, similar to what is seen in the RNA-Seq of quiescent and activated MuSCs (Figure 3E). Together, the data indicates that the Nedd4L-cKO MuSCs exhibit a gene signature suggestive of activated MuSCs.

### Nedd4L-cKO MuSCs are transcriptionally similar to active MuSCs

The RNA-Seq data suggest that the Nedd4L-cKO MuSCs are no longer in a quiescent state. However, when we compare the expression of the myogenic regulatory factors (MRFs), we see that there is no difference in the expression of any of the MRFs, and only a slight non-significant decrease in *Pax7* expression, possibly due to its reduced mRNA stability in the Nedd4L-cKO (Supplemental Figure 2D, Supplemental Figure 4A-E). Further, there is no significant increase in *Ki67* expression in the Nedd4L-cKO cells (Supplemental Figure 4F). Together, this would indicate that MuSCs from the Nedd4L-cKO mouse have not fully activated and are not proliferating. Therefore, we posit that the genetic deletion of Nedd4L leads to MuSCs entering a state of G_alert_^6^, where the cells have not broken quiescence, but are closer to activation than a fully quiescent WT MuSC.

To confirm this, we compared the changes in the transcriptome of WT vs Nedd4L-cKO MuSCs to the changes in the transcriptome of quiescent (QSC) vs activated (ASC) MuSCs (Figure 4A). The QSC vs ASC data was retrieved from publicly available data (GSE144871)^11^. The QSC cells are from uninjured muscle while the ASC are from the tibialis anterior (TA) and gastrocnemius muscles 3 days post CTX induced injury. We see from this comparison that the changes in the transcriptome between WT and Nedd4L-cKO is similar to the changes that occur during injury induced activation (Figure 4A).

**Figure 4:**
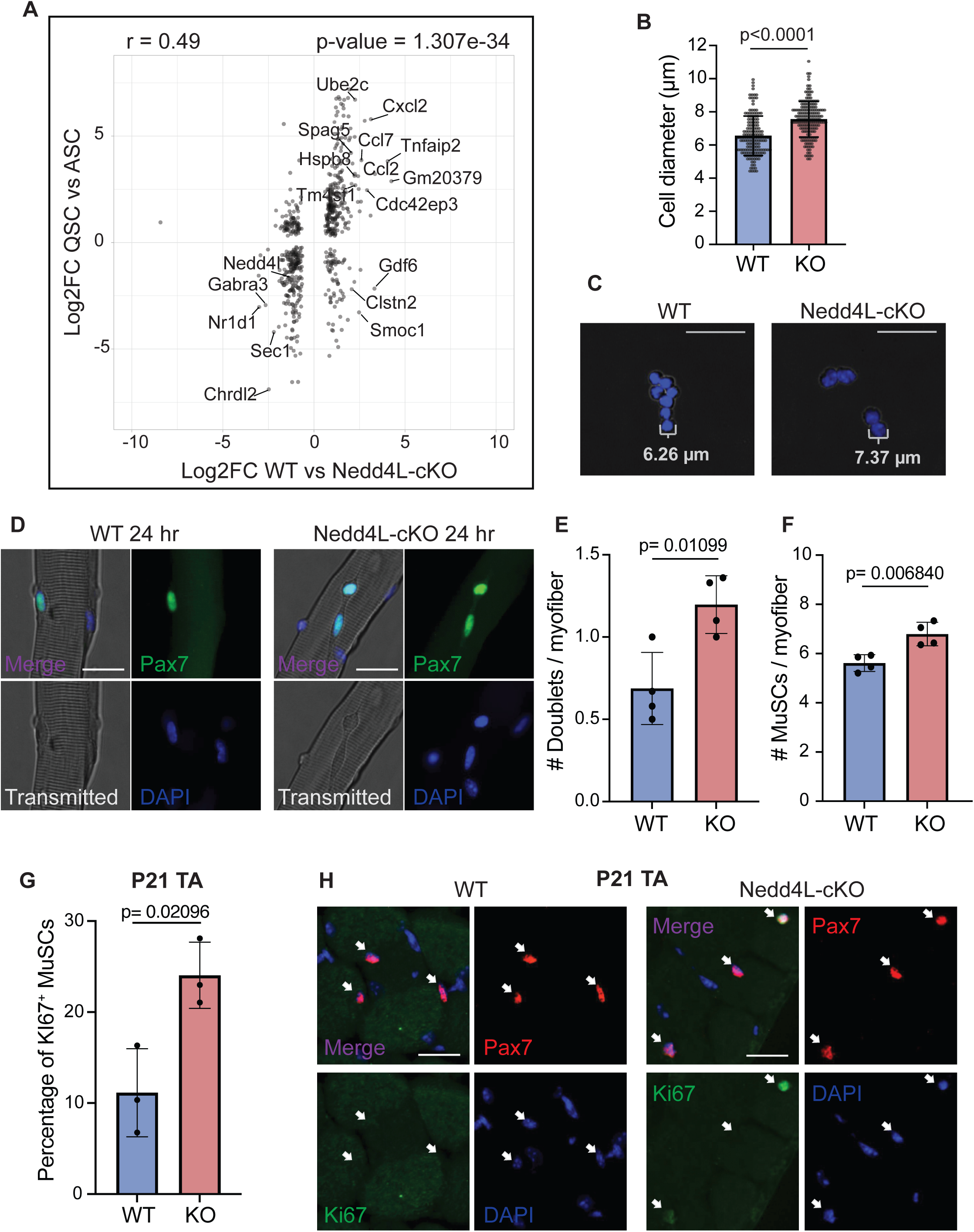
Nedd4L-cKO MuSCs are transcriptionally similar to active MuSCs. **A**. Scatterplot comparing the LFC of WT vs Nedd4L-cKO with the LFC of quiescent muscle stem cells (QSC) vs activated muscle stem cells (ASC). The QSC and ASC sequencing data is publicly available and retrieved from GSE144871^11^. Genes with a mean expression of over 200 normalized counts and an adjusted p-value <0.1 were plotted. Genes with an absolute LFC >2 in both conditions were labelled, with Nedd4L being manually labelled. **B**. Bar graph of the cell size of freshly isolated MuSCs from WT and Nedd4L-cKO mice, n = 3 mice, 163 WT cells and 184 KO cells. **C.** Representative images of freshly isolated MuSCs. **D.** Representative images of EDL myofibers 24 hours post isolation stained for Pax7, scale bar = 25 µm. **E.** Bar graph of the number of MuSC doublets per myofiber, n = 4 mice. **F.** Bar graph of the number MuSCs per myofiber, n = 4 mice. **G.** Bar graph representing the percentage of Ki67^+^ MuSCs from WT and Nedd4L-cKO P21 TA muscle, n = 3 mice. **H.** Representative images of TA cross sections from P21 WT and Nedd4L-cKO mice, stained for Pax7 and Ki67, scale bar = 25 µm. Data presented as mean ± SD.

The state of G_alert_ is affiliated with an increase in cell size and a shorter time to cell division^6^. We see in freshly sorted MuSCs that the Nedd4L-cKO condition has significantly larger cells (Figure 4B-C). Additionally, in EDL myofibers cultured for 24 hours, we found that there are significantly more MuSC doublets and a higher number of MuSCs in the Nedd4L-cKO (Figure 4D-F). This would imply that the Nedd4L-cKO MuSCs more readily divide when cultured compared to their WT counterparts. Further, in the TA cross sections from P21 (post-natal day 21) mice, we see that the Nedd4L-cKO MuSCs have a higher proportion of those that are Ki67 positive (Figure 4G-H). At 3 weeks of age, MuSCs will enter quiescence for the first time^52^. This higher percentage of Ki67 cells suggests that the genetic deletion of Nedd4L impedes the ability of the MuSCs to acquire quiescence. Together, the data presented here strengthens the hypothesis that Nedd4L maintains MuSC quiescence and with its genetic deletion the cells enter G_alert_.

### Genetic deletion of Nedd4L alters the proteome of freshly isolated muscle stem cells

Nedd4L is a member of the UPS system and as such regulates the proteome of a cell. In order to determine how the loss of Nedd4L affects the proteins in quiescent muscle stem cells we performed Label Free Quantification (LFQ) of proteins with nano LC-MS/MS (Figure 5A). Every sample consisted of 200,000 MuSCs, necessitating the pooling of multiple mice.

**Figure 5:**
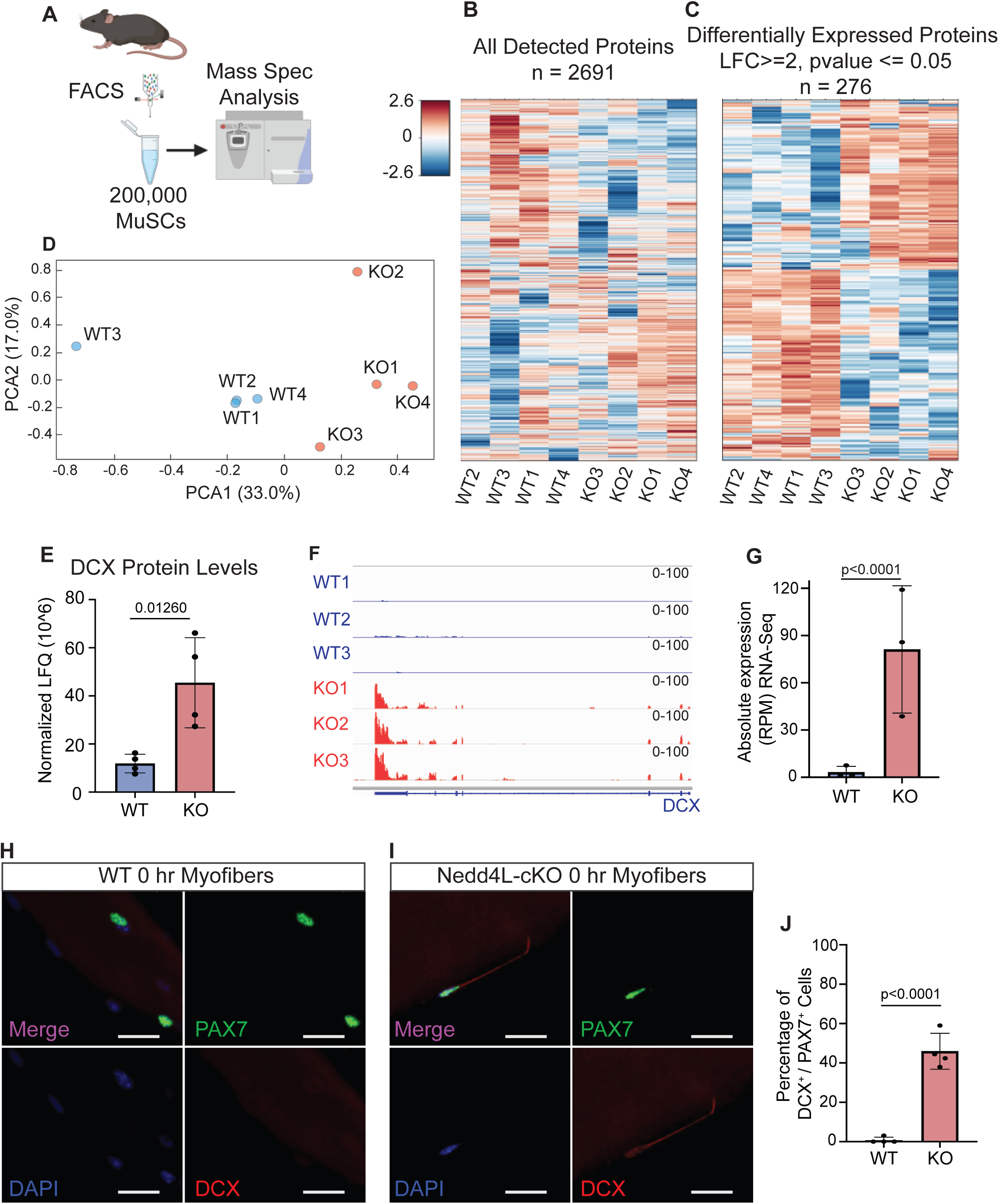
Genetic deletion of Nedd4L alters the proteome of freshly isolated muscle stem cells. **A.** Diagram of the design of the Mass spectrometry experiment performed on 200,000 freshly isolated WT and Nedd4L-cKO MuSCs, generated with Biorender.com. **B.** Heatmap of all detected proteins in 200,000 freshly isolated MuSCs from WT and Nedd4L-cKO mice. **C.** Heatmap of the protein levels in WT and Nedd4L-cKO MuSCs for all significantly different proteins (LFC ≥ 2, p-value ≤ 0.05). **D**. PCA plot of WT and Nedd4L-cKO MuSCs. **E**. Bar graph representing the DCX protein levels from the Mass spec. **F.** IGV tracks of RNA-Seq reads for Dcx. **G**. Bar graph of the absolute expression (RPM) of Dcx in WT and Nedd4L-cKO mice. **H**. Immunofluorescence for DCX and PAX7 in WT MuSCs associated with freshly isolated (0hr) EDL myofibers. **I**. Immunofluorescence for DCX and PAX7 in Nedd4L-cKO MuSCs associated with freshly isolated EDL myofibers. **J**. Bar graph representing the percentage of Dcx^+^/Pax7^+^ MuSCs associated to freshly isolated EDL myofibers. Data presented as mean ± SD.

When we analyzed the proteome of freshly isolated WT and Nedd4L-cKO MuSCs we were able to detect 2691 proteins and found that 276 of them were significantly different between the conditions (LFC ≥ 2, p-value ≤ 0.05) (Figure 5 B-C). We observed that this caused a distinct proteomic profile between the WT and Nedd4L-cKO MuSCs as seen by the separate clustering on a PCA plot (Figure 5D). We confirm with the Mass spec data the loss of CHRDL2 protein in the Nedd4L-cKO MuSCs (Supplemental Figure 5A).

A particularly interesting protein that was detected in the Nedd4L-cKO cells is Doublecortin (DCX) (Figure 5E). DCX is a 40 kDa protein and is most well-known for its role in neuronal migration and neurogenesis^53–55^. This protein has also been shown to have a role in MuSCs; through sequencing it was demonstrated that the gene is only expressed in the cases of neonatal MuSCs, activation, and in the dystrophic (mdx) mouse model^56^. Further, it was demonstrated to have a key role in the differentiation of MuSCs and their fusion to pre-existing fibers during regeneration^57^. This lead us to compare the transcriptomic expression of *Dcx* from our RNA-Seq data and found that the gene is significantly upregulated in the Nedd4L-cKO MuSCs, while having almost no detectable expression in the WT MuSCs (Figure 5F-G). In order to validate the expression of DCX protein in quiescent MuSCs, we stained MuSCs associated to freshly isolated EDL myofibers, for PAX7 and DCX. From this we observed that while almost no WT MuSCs were positive for DCX protein, a large percentage of Nedd4L-cKO MuSCs were positive, approximately 45% (Figure 5H-I). This level of DCX positive MuSCs was maintained in older mice, aged 6 months (Supplemental Figure 5 B-D). Together, we posit that the genetic deletion of Nedd4L leads to a change in the proteome, resulting in the MuSCs entering G_alert_ and being primed to differentiate and repair damaged myofibers. Further the expression of DCX is a marker for this intermediary stage of MuSC activation.

### Chromatin accessibility is unaltered by the genetic deletion of Nedd4L

Breaking quiescence and activation is a complex process that results in numerous transcriptomic and biochemical changes in MuSCs, along with an alteration in the chromatin state^5, 14^. We sought to determine how the chromatin of these cells that were shifted closer to activation were affected. To accomplish this, we performed Assay for Transposase Accessible Chromatin (ATAC-Seq) on freshly isolated WT and Nedd4L-cKO MuSCs (Figure 6A).

**Figure 6:**
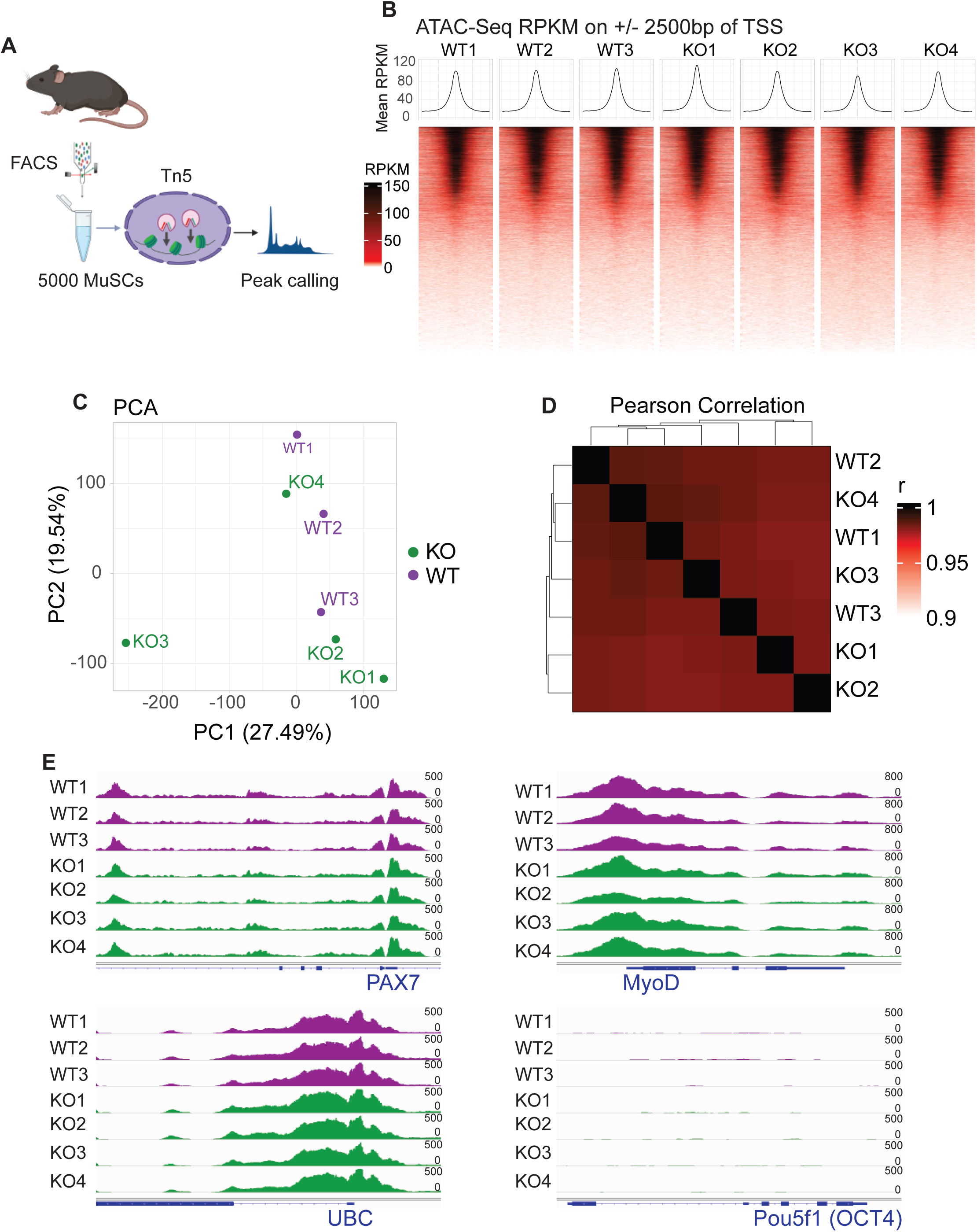
Chromatin accessibility is unaltered by the genetic deletion of Nedd4L. **A**. Schematic of the ATAC-seq experiment performed on 5000 freshly isolated MuSCs, generated with Biorender.com. **B**. Pile up heatmap of the ± 2500 bp region around TSS of all genes in each ATAC-seq sample. **C**. PCA plot of WT and Nedd4L-cKO MuSC ATAC-seq samples. **D.** Heatmap of the sample-to-sample Pearson correlation of WT and Nedd4L-cKO MuSC ATAC-seq samples. **E**. IGV tracks of ATAC-Seq reads for myogenic genes Pax7 and MyoD, housekeeping gene ubiquitin (UBC), and Pou5f1 (OCT4) as a negative control.

Interestingly, we found that there is no difference in chromatin accessibility between the WT and Nedd4L-cKO MuSCs. We verified the quality of our ATAC-seq and found that the results, including the pile-up of reads around TSS and peak annotations, showed a typical profile expected from ATAC-Seq (Figure 6B). We see that there is no clustering in the PCA, nor any difference in the Pearson correlation between the samples, regardless of their genotype (Figure 6C-D). As a positive control, we visualized PAX7, MYOD1 and the housekeeping gene UBC and saw no differences between samples in the accessibility at the promotor of these genes (Figure 6E). Finally, as a negative control the promoter of *Pou5f1* (*OCT4*) was inaccessible as expected (Figure 6E). When we looked at individual genes that were identified from our RNA-Seq as having a change in their expression, we again see that there is no difference in chromatin accessibility (Supplemental Figure 6).

Together, this data indicates that the transition from a deeper state of quiescence (PAX7^+^/DCX^-^) to the PAX7^+^/DCX^+^ state does not lead to any changes in chromatin accessibility.

### Nedd4L-cKO muscle stem cells are more prone to differentiation

To determine the functional effect of the genetic deletion of Nedd4L in MuSCs, EDL myofibers were isolated from 6-8 weeks old male and female mice, and cultured for 0, 24, 48 and 72 hours. When stained for PAX7 we found that there was no difference in the number of MuSCs in freshly isolated myofibers from Nedd4L-cKO and WT mice (Figure 7 A,B). However, as they grew in culture for 48 and 72 hours, it was observed that the Nedd4L-cKO MuSCs could not expand as rapidly as the WT, as seen by the reduced number of PAX7^+^ cells in the Nedd4L-cKO condition (Figure 7 C-F). To validate that this was not due to apoptosis, myofibers that were cultured for 72 hours were stained for cleaved Caspase-3, a marker of apoptosis (Supplemental Figure 7A-B). We found no difference in the percentage of MuSCs that were apoptotic between the WT and Nedd4L-cKO.

**Figure 7:**
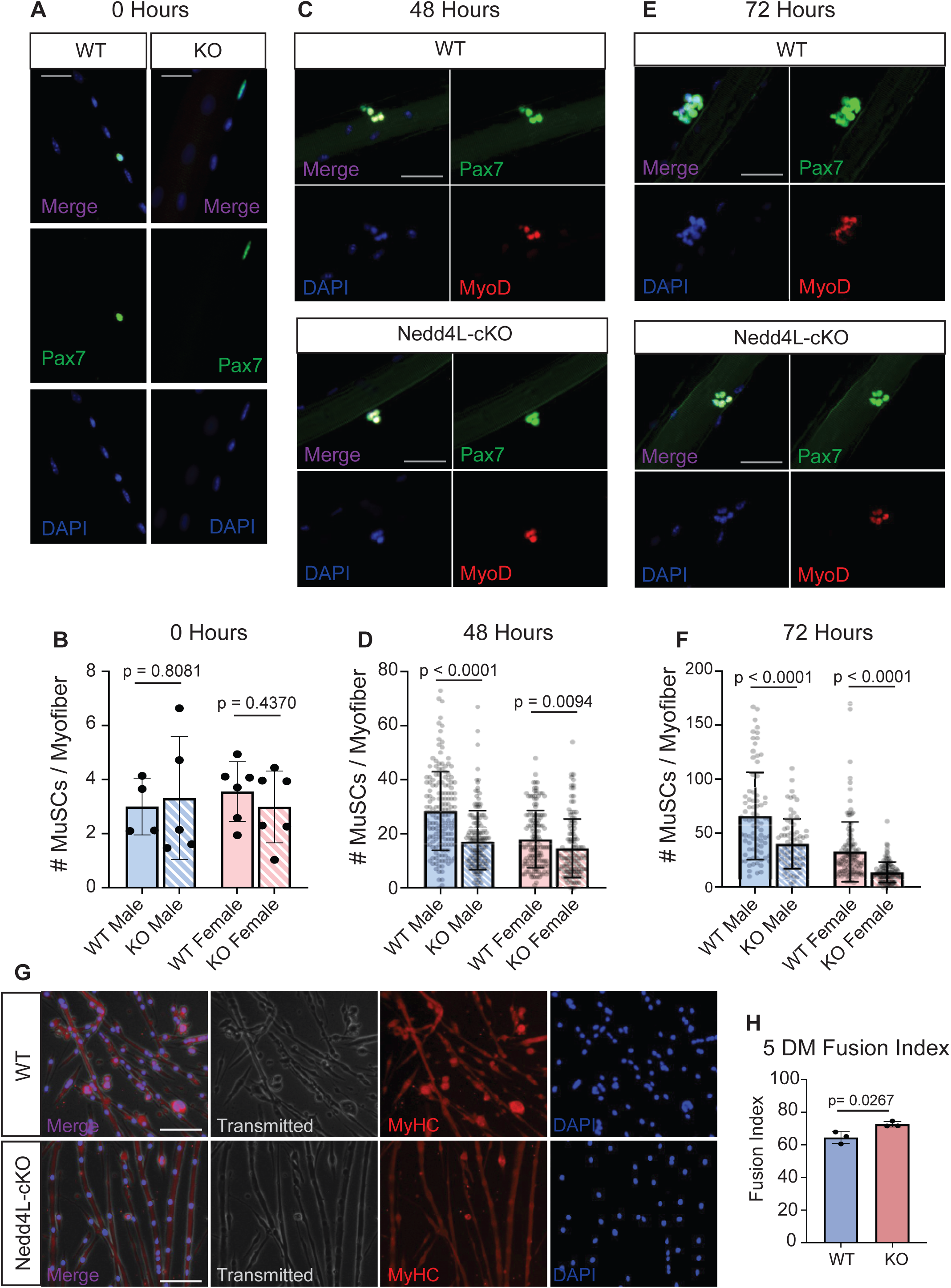
Nedd4L-cKO muscle stem cells are more prone to differentiation. **A.** Representative images of isolated EDL myofibers 0 hours post isolation, stained for PAX7, scale bar = 25 µm. **B.** Bar graph representing the number of MuSCs per myofiber 0 hours post isolation in male and female WT and Nedd4L-cKO mice, aged 6-8 weeks. n = 4-6 mice. **C.** Representative images of isolated EDL myofibers 48 hours post isolation, stained for PAX7 and MyoD1, scale bar = 25 µm. **D.** Bar graph representing the number of MuSCs per myofiber 48 hours post isolation in male and female WT and Nedd4L-cKO mice, aged 6-8 weeks. n = 4-6 mice. **E.** Representative images of isolated EDL myofibers 72 hours post isolation, stained for PAX7 and MyoD1, scale bar = 25 µm. **F.** Bar graph representing the number of MuSCs per myofiber 72 hours post isolation, in male and female WT and Nedd4L-cKO mice, aged 6-8 weeks. n = 4-6 mice. **G.** Representative images of WT and Nedd4L-cKO primary myotubes grown in differentiation media for 5 days and stained for MyHC. **H.** Bar graph representing the fusion index of WT and Nedd4L-cKO grown for 5 days in differentiation media. Data presented as mean ± SD.

The inability of the Nedd4L-cKO MuSCs to properly expand may be due to defects in proliferation. To determine this, we performed an EdU assay on cultured WT and KO primary myoblasts for 12 and 24 hours. We see that there is no difference in the number of EDU positive cells between the conditions, thereby indicating that there is no difference in proliferation upon genetic deletion of Nedd4L (Supplemental Figure 7C-D).

An increase in the level of differentiation could also explain the lower number of MuSCs. When performing a differentiation assay for 5 days, we observed that the fusion index in the Nedd4L-cKO myotubes was significantly higher than in the WT, indicating an increase in differentiation (Figure 7 G-H). As previously mentioned from our RNA-Seq data, there is also large number of genes involved in late differentiation that are upregulated in the Nedd4L-cKO MuSCs. Together, this data suggests that the loss of Nedd4L pushes the MuSCs towards precocious differentiation, thereby hindering the expansion of the MuSC pool. However, in cultured primary myoblasts, we see no difference in the protein level of MyoD or Pax7, nor do we see any difference in the protein level of Myogenin in 5DM cultured myotubes (Supplemental Figure 7E-F). Interestingly, while we do not see a change in MuSC numbers in EDL myofibers at T0 in young mice, we do see a decrease in the EDLs of 6 month old Nedd4L-cKO mice compared to WTs of the same age (Supplemental Figure 7G-H). This could be indicative of a gradual decline in MuSC number due to precocious differentiation as the mice age.

## Discussion

Adult stem cell quiescence is a rather dynamic state, with the cells continuously monitoring and responding to a plethora of signals by modulating its depth of quiescence^5, 44, 58, 59^. Quiescence should be viewed as a gradient, rather than an on/off switch, with the cells able to move from a state of deep quiescence to one viewed as G_alert_, passing through a number of checkpoints. G_alert_ may itself possess a gradient ranging from more quiescent and more active.

Previous studies have shown the importance of proteostasis for MuSC function and quiescence^26, 60, 61^. In this study we show that Pax7 directly regulates MuSC quiescence via its transcriptional control of the E3 ubiquitin ligase Nedd4L.

When Nedd4L is deleted in MuSCs, there is an alteration in the transcriptome with an upregulation of genes involved in MuSC activation and differentiation. Notably, we observed the loss of expression of *Nr1d1*, *Calcr* and *Chrdl2* in the Nedd4L-cKO MuSCs, all of which have been shown to play a role in the maintenance of quiescence^39–43^. Interestingly, while there is an upregulation of genes involved in myogenic differentiation, most of them are characteristic of late differentiation (Supplemental Figure 3). Of note, *Mymk* is upregulated in the Nedd4L-cKO MuSCs, which is essential for the fusion of myoblasts, a process that is important for the late stages of differentiation^62, 63^. Despite these transcriptomic changes, none of the MRFs are deregulated, nor do we observe an increase in *Ki67* expression in the Nedd4L-cKO MuSCs. This lead us to conclude that the loss of Nedd4L results in the cells entering G_alert_, as they are less quiescent than WT cells, exhibiting a larger cell size and upregulation mitochondrial genes, but they have not yet re-entered the cell cycle.

Furthermore, we identify a new marker for the intermediary state between quiescence and activation, PAX7^+^/DCX^+^/MYOD1^-^, as we see that only the Nedd4L-cKO MuSCs would consistently express DCX with no change in Myod1 expression. Previous work on this protein has shown that, in MuSCs, DCX is only expressed in the *in vivo* context, seen only during injury, neonatal mice or in mdx mice, but not in cultured myoblasts^56, 64^. A previous study by Ogawa et al. demonstrated that DCX is involved in the differentiation of MuSCs, specifically with regards to the fusion of MuSCs to pre-existing myofibers, in a manner that does not follow the classic myogenic differentiation pathway^64^. This finding could explain why there is a large number of late differentiation genes that are upregulated in the Nedd4L-cKO cells, with few of the genes associated with early-stage differentiation exhibiting any significant change in their expression. Further, this push towards differentiation and fusion with existing fibers would explain the difficulty the Nedd4L-cKO MuSCs have in expanding when the myofibers are cultured. This is despite not having any observed defect in proliferation, nor an increase in the apoptosis marker cleaved Caspase-3.

An accepted role for Nedd4L is that of a tumor suppressor, it is most often downregulated in cancer cells^27, 33–36^. This role would explain its function in the maintenance of MuSC quiescence. We hypothesize that in quiescent MuSCs, Nedd4L would act as a tumor suppressor and inhibit the MuSCs ability to enter the cell cycle.

Our results demonstrate that protein turnover, mediated by the UPS, is a major factor in the maintenance of MuSC quiescence. We propose that MuSCs continuously produce proteins necessary for their activation, while simultaneously degrading them. The rationale for this phenomenon is that injury is completely unpredictable, both in timing and in intensity. MuSCs must therefore always be primed to enter the cell cycle in order to rapidly respond to these unexpected injuries and regenerate the tissue as effectively as possible. Therefore, active recycling of protein involved in early stages of MuSCs activation can be viewed as a cell survival premium to ensure against the unpredictable nature of tissue injury. Our study indicates that PAX7 regulation of Nedd4l via recruitment of UPS system serves to maintain MuSCs in quiescent state yet poised for rapid entry into cell cycle. Further research will need to be conducted to fully explore the role of the UPS in muscle stem cell function and on the role protein turnover plays in stem cell quiescence.

**Table 1:**
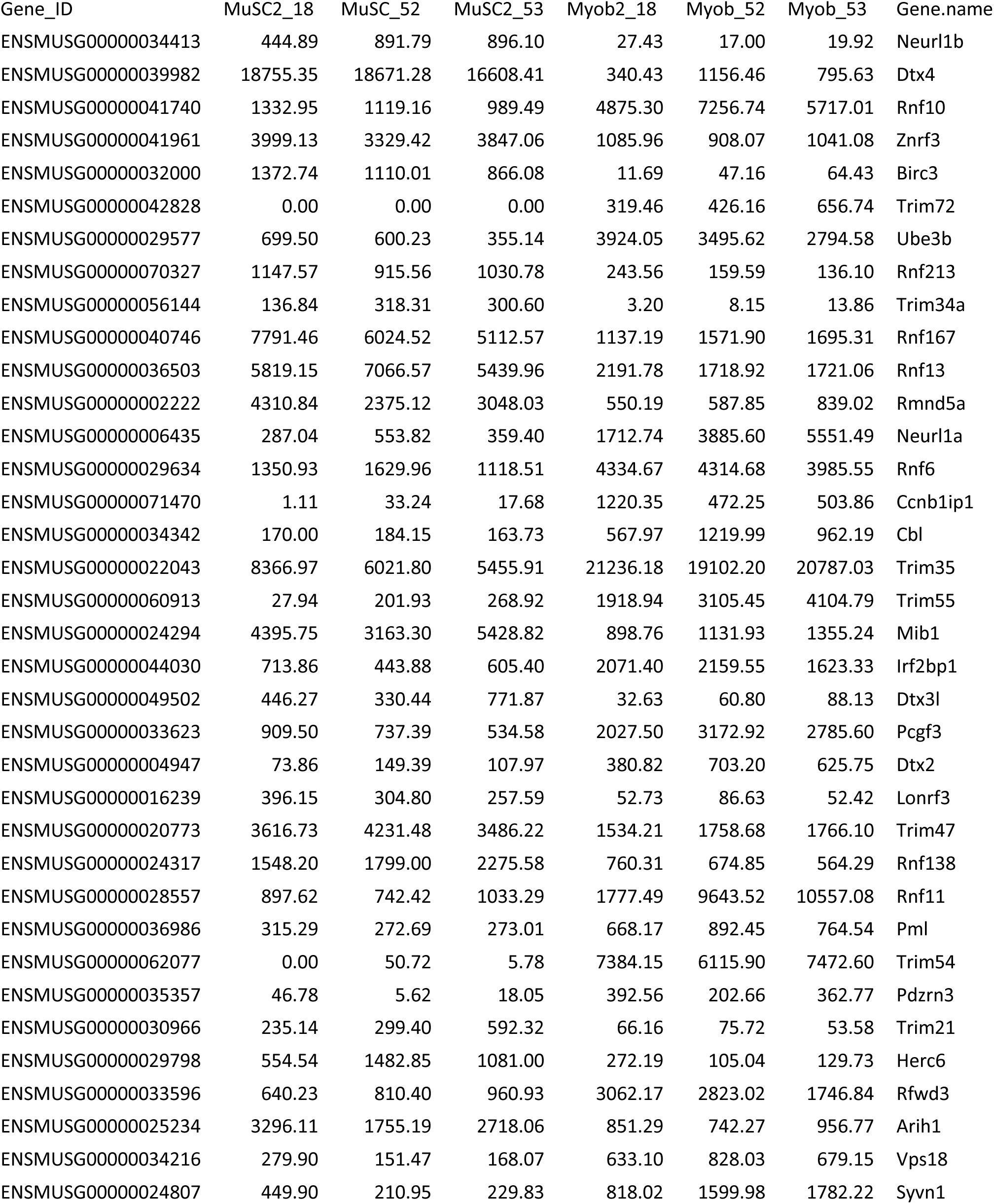
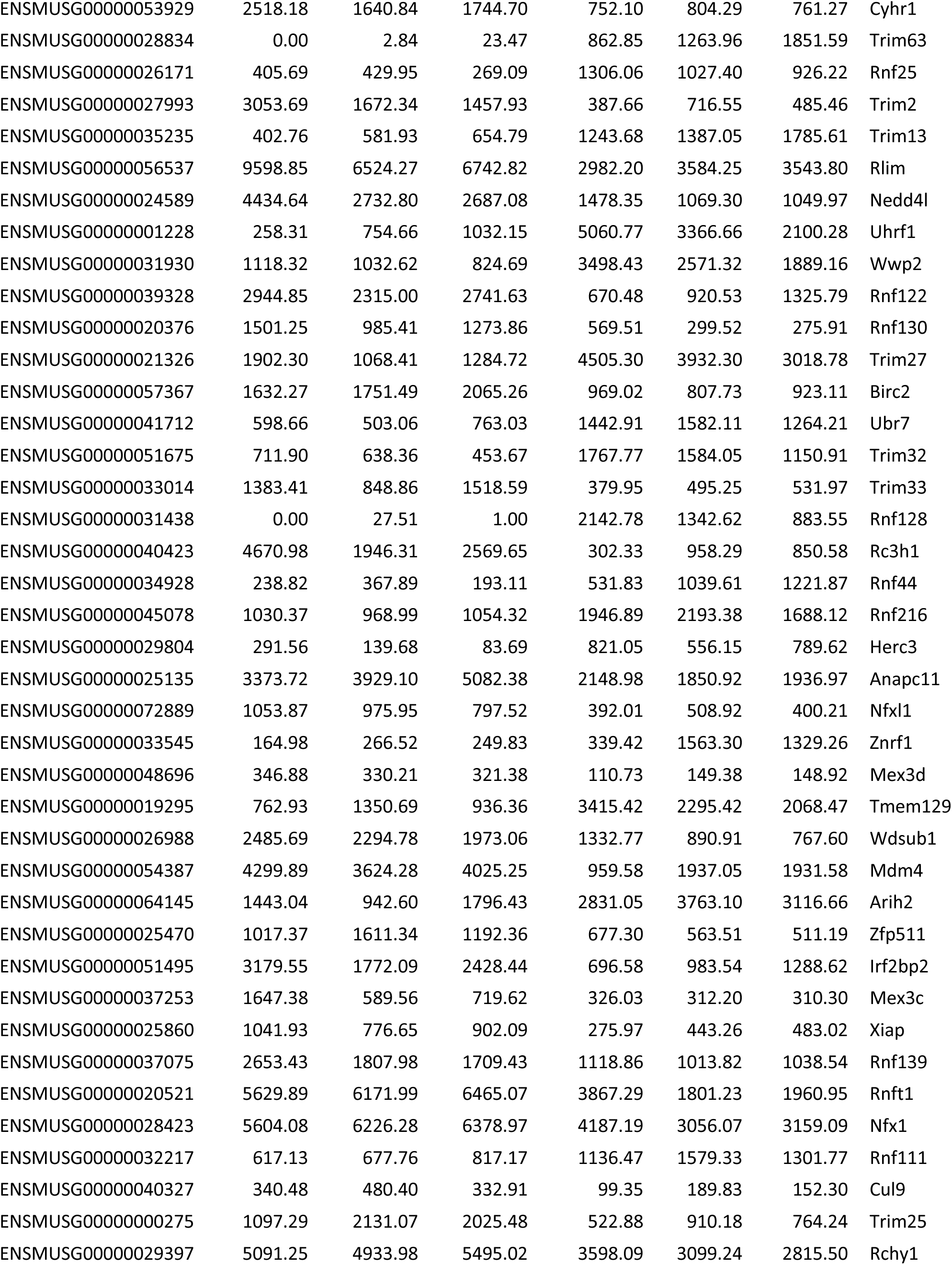
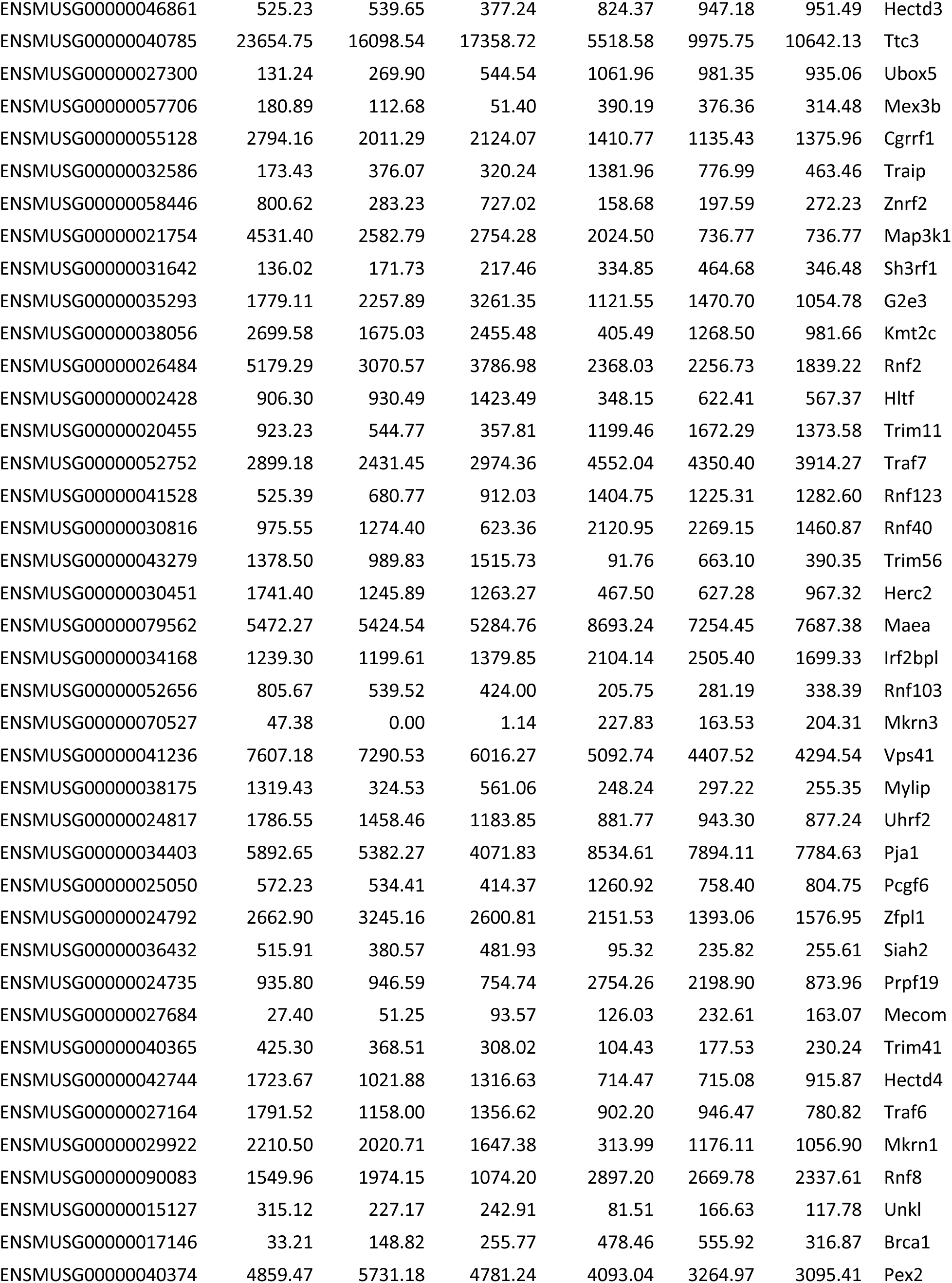
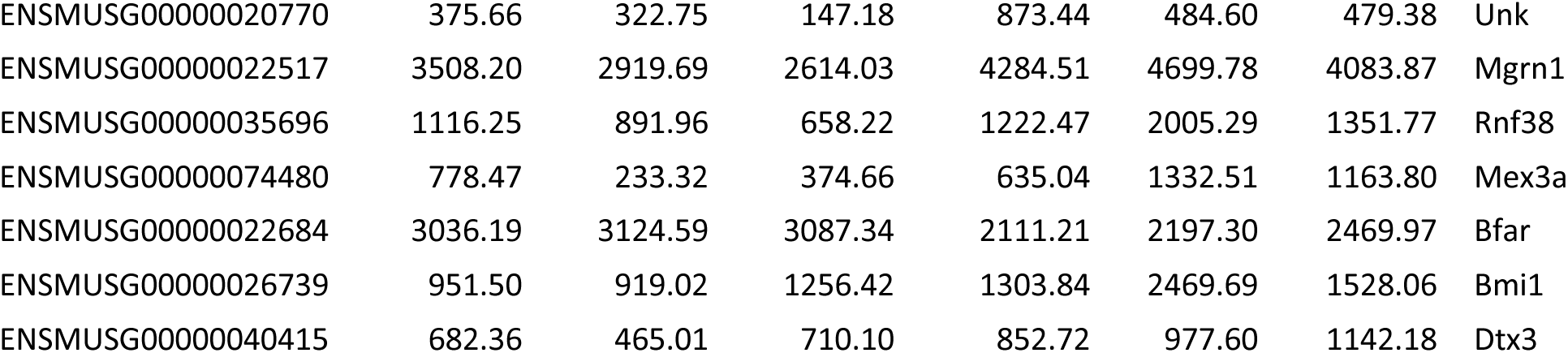
List of E3 ubiquitin ligases expressed at a minimum 100 base counts from the RNA-Seq of muscle stem cells and/or primary myoblasts

## Acknowledgments

We thank Christian Young and Mathew Duguay at the Lady Davis Institute for Medical Research— Jewish General Hospital—core facility for help with fluorescence-activated cell sorting (FACS). Research in Soleimani laboratory is supported by the Natural Sciences and Engineering Research Council (NSERC), RGPIN-06774 and by Project grant PJT-156087 from the Canadian Institute of Health Research (CIHR). D.M.B and K.S. are supported by David G. Guthrie Fellowship in Medicine from McGill University. CHB is grateful to Genome Canada for financial support through the Genomics Technology Platform (GTP) (264PRO), and for support from the Segal McGill Chair in Molecular Oncology at McGill University (Montreal, Quebec, Canada). CHB is also grateful for support from the Warren Y. Soper Charitable Trust, the Alvin Segal Family Foundation of the Jewish General Hospital (Montreal, Quebec, Canada), and the Terry Fox Research Institute.

## Author Contributions

Conceptualization, V.D.S.; Methodology, D.M.B., K.S., H.K., A.H.C., H.S.N., and V.D.S.; Formal Analysis, D.M.B., V.D.S, A.H.C., G.P., V.R., R.Z., H.S.N.; Investigation, D.M.B., V.D.S., K.S., F.L., L.R., V.R.; Resources, V.D.S, H.K.; Writing – Original Draft, D.M.B., and V.D.S.; Writing – Review & Editing, D.M.B., V.D.S., H.S.N and A.J.; Visualization, D.M.B. and V.D.S.; Supervision, V.D.S., A.J., C.B., H.S.N.; Funding Acquisition, V.D.S.

## Methods

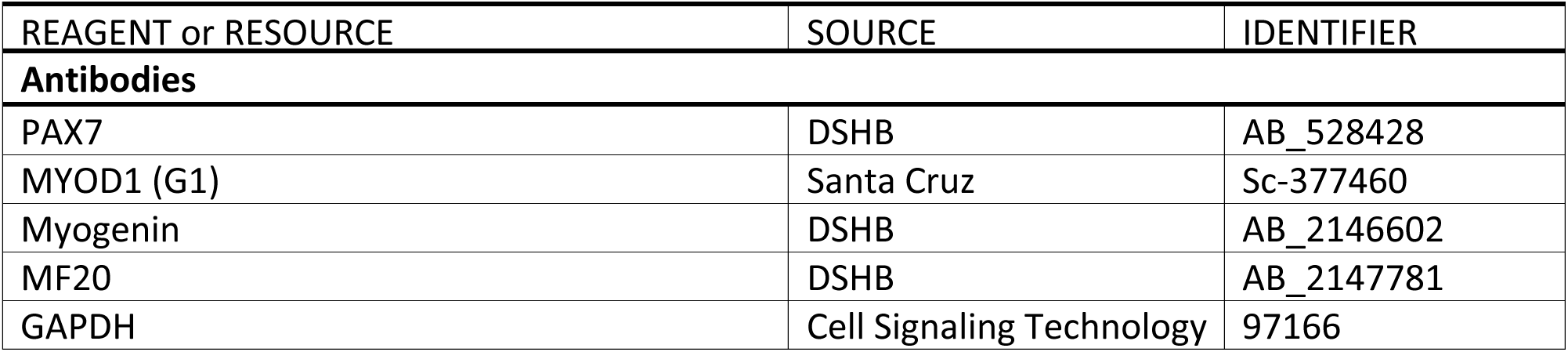
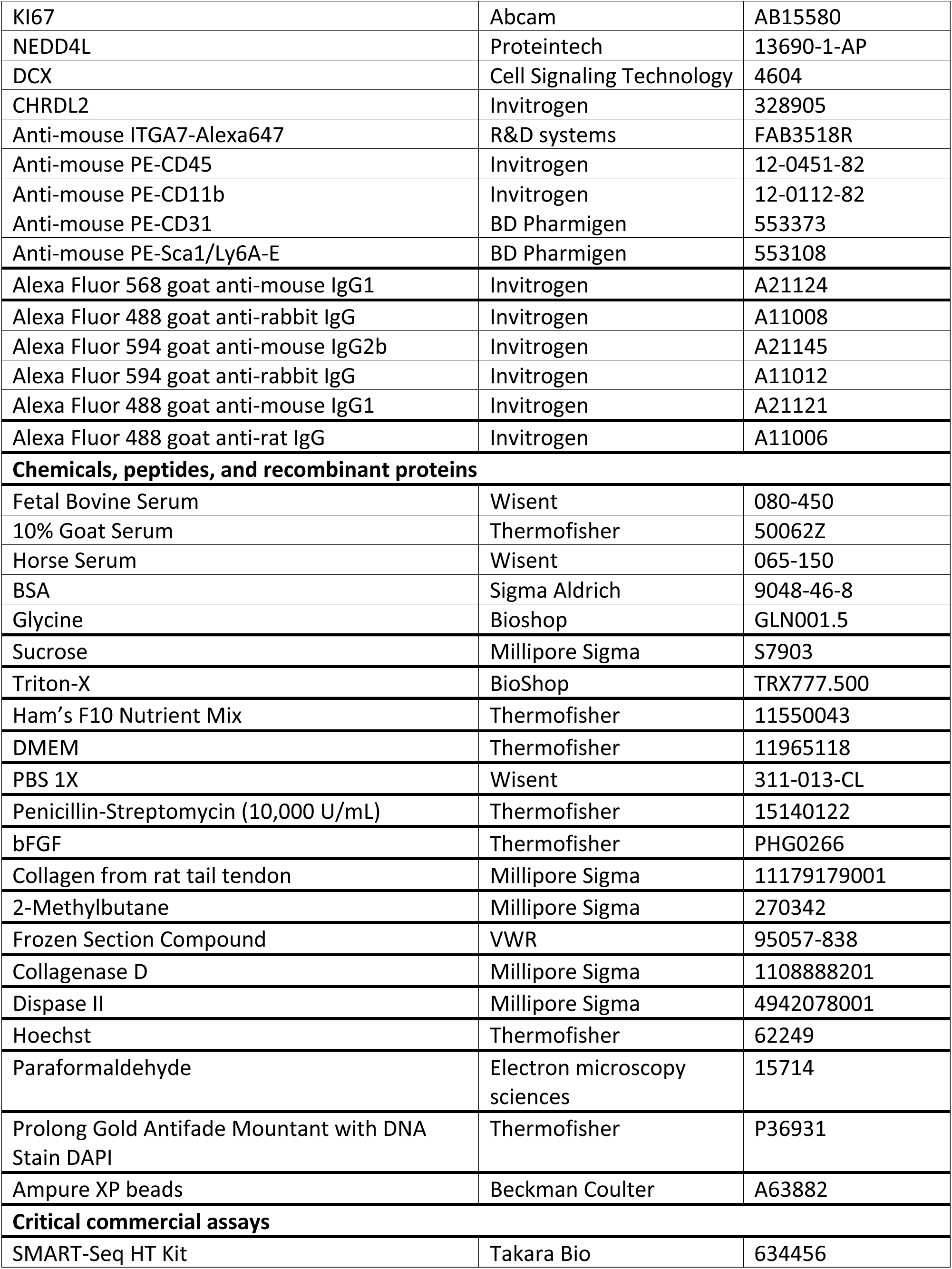
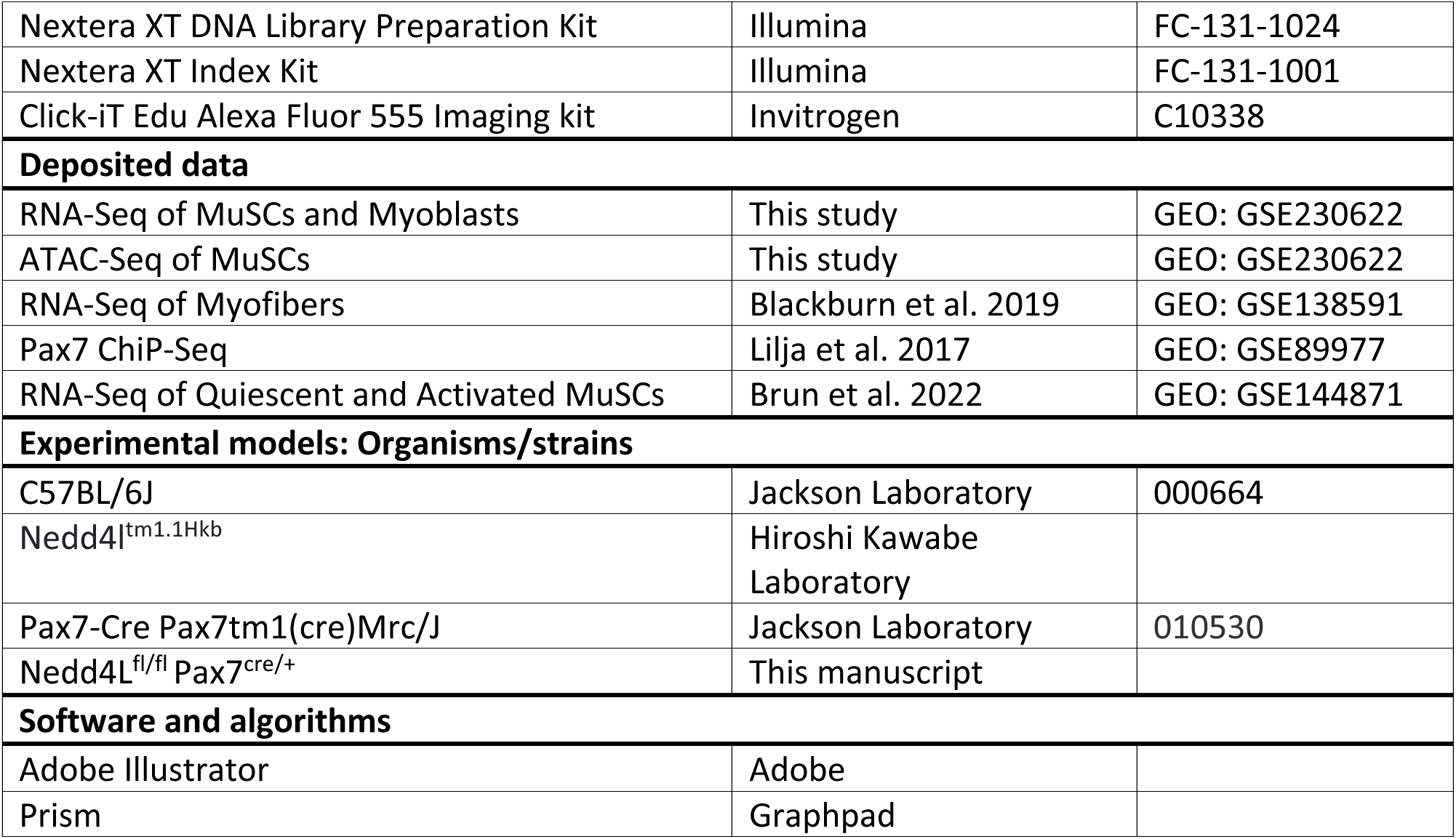

### Mouse strains and animal care

All animal procedures in this study for mice were approved and performed in accordance with the guidelines of the McGill University Animal Care Committee (UACC). The mouse line used in this study is Nedd4L^fl/fl^ Pax7^cre/+^ with a C57BL/6 background. In all experiments, mice are age and sex matched.

### Intramuscular injection of MG132 into the tibialis anterior

8 week of C57BL/6 mice were injected intramuscularly with 50 µL of 10mM MG132 dissolved in DMSO, once every 24 hours for 3 days. Control mice were injected with 50 µL of DMSO for the same time period. After the final injection, TAs were collected 24 hours later and fixed in 0.5% PFA for 2 hours at 4°C. The TAs were placed in 20% sucrose at 4°C overnight and then frozen in frozen section compound using liquid nitrogen cooled 2-methylbutane.

### Immunofluorescence of TA muscles

When the TA muscles are dissected from the mice, they are placed in a 1.5 mL tube and are fixed immediately in 1 mL of 0.5% PFA in PBS for 2 hours at 4°C. The TA is then incubated in 20% sucrose in dH_2_O at 4°C overnight. The TA muscle was frozen in clear sectioning liquid using liquid nitrogen cooled isopentane.

The TAs were sectioned at a width of 9 µm using a cryostat at −25°C and placed on a slide. A hydrophobic barrier was drawn around the cross sections using a pap pen. The slide was rinsed with PBS and the cross sections were then permeabilized with 0.5% Triton-X for 15 minutes at RT. The sections were then washed 3 times with PBS and blocked with blocking buffer (10% goat serum, 3% BSA, 0.1M glycine) for 2 hours at RT. Primary antibodies were diluted in the blocking buffer and the cross sections were incubated overnight at 4°C in a humid chamber.

The next day, cross sections were washed in 0.1% Triton-X in PBS 3 times at RT. Secondary antibodies were diluted 1:400 in blocking buffer and cross sections were incubated for 1 hour at RT. Slides were then washed 3 times with 0.1% Triton-X in PBS. Slides were mounted using prolong gold antifade with DAPI.

### Fluorescence activated cell sorting of muscle stem cells

The hindlimb muscles were dissected and minced using scissors and subsequently digested in 5 mL of FACS digestion media (HAM’s F10 with 2.4 U/mL Collagenase D, 10 U/mL Dispase II, and 0.5 mM CaCl_2_) for 30 minutes at 37°C in a 5% CO_2_ tissue culture incubator. The muscle was pulsed in a centrifuge and the supernatant was transferred to 8 mL of FBS on ice. The remaining muscle pellet is triturated with a pipette and 5 mL of FACS digestion media is added and the muscle was incubated for another 30 minutes. The digested muscle was transferred to the same FBS. Sample was then filtered through a 40 µm cell strainer. The cells are pelleted by centrifugation at 500 G for 18 minutes at 4°C. The cell pellet was resuspended in 500 µL of 2% FBS and 0.5mM EDTA in PBS and were incubated for 20 minutes at room temperature with the following antibodies: Alexa647-conjugated to anti-mouse ITGA7, PE-CD31, PE-CD45, PE-CD11b, PE-SCA1 and Hoechst 33342. After the cells were washed with 10 mL of 2% FBS in PBS and centrifuged at 500 G for 15 minutes at 4°C. Cells were resuspended in 800 µL of 2% FBS in PBS and filtered through a 40 µm cell strainer. The ITGA7+/Hoechst+/CD31−/CD45−/CD11b−/SCA1− population was sorted with a FACS aria fusion, as previously described^65^.

### Cell culture

Muscle stem cells that were isolated by FACS were cultured on collagen coated plates in growth media (Ham’s F10 supplemented with 20% FBS, 1% penicillin/streptomycin and 5 ng/mL of bFGF) in a cell incubator set to 37°C and 5% CO_2_. Cells were passaged upon attaining 70% confluence.

### Creating PAX7 or NEDD4L over-expressing primary myoblasts

Stable cell lines of primary myoblasts over-expressing PAX7 or NEDD4L were generated as previously described^66^. Briefly, retrovirus particles containing the plasmids of interest were generated in Phoenix helper-free retrovirus producer lines with lipofectamine. The viral supernatant was collected 48 h after transfection and subsequently applied to cultured primary myoblasts for 8 h. Afterwards the cells were washed twice with 1 × PBS and grown for 48 h in normal growth media. Puromycin selection media (HAM’s F10 supplemented with 20% FBS, 5 ng/ml bFGF, 1% penicillin/streptomycin, and 2.5 μg/ml puromycin) was then given to the cells for 1 week. Finally, the cells were maintained in low puromycin maintenance media (HAM’s F10 supplemented with 20% FBS, 5 ng/ml bFGF, 1% penicillin/streptomycin, and 1.25 μg/ml puromycin).

### EdU incorporation assay

Cultured primary myoblasts were treated with EdU as described in the Click-It Plus EdU Alexa Fluor 555 Imaging Kit (Invitrogen C10638). Briefly, growth media with 10 µM of EdU was added to primary myoblasts seeded on collagen coated chamber slides. The cells were then cultured for 12 or 24 hours. Media was removed and the cells washed twice with PBS and then fixed with 3.2% paraformaldehyde. EdU staining was performed as described in the kit.

### Myoblast differentiation

Cultured myoblasts were grown to 90% confluence and differentiated in DMEM supplemented with 5% horse serum (HS) and 1% penicillin/streptomycin for 5 days. After 3 days the media was replaced with fresh differentiation media for the duration of the assay.

### Preparation of SMART RNA-Seq libraries from freshly sorted muscle stem cells

1000 MuSCs were FACS sorted directly into 10 µL SMART lysis buffer (1 µL SMART-Seq Reaction Buffer (95% SMART-Seq 10X lysis buffer, 5% RNAse Inhibitor), 9 µL ddH_2_O) in a 0.2 mL microtube. 1 µL of 3’ SMART-Seq CDS primer II A was added, and the samples were incubated for 3 minutes at 72°C, then placed on ice. 12.5 µL of the template switching master mix (0.7 µL of nuclease free water, 8 µL of one-step buffer, 1 µL SMART-Seq HT oligonucleotide, 0.5 µL RNase inhibitors, 0.3 µL of SeqAMP DNA polymerase and 2 µL of SMARTscribe reverse transcriptase) was added to each sample. The cDNA was amplified for 11 cycles as described previously^67, 68^.

The cDNA was purified using AMPure beads at a 1:1 volume/volume ratio. The beads were washed twice with 200 µL of 80% ethanol. The cDNA was eluted from the beads with 17 µL of TE buffer (10 mM Tris-HCl, 1mM EDTA, pH 8.0). 250 pg of cDNA in a final volume of 1.25 µL, was transferred to a fresh microtube. With the Nextera XT DNA Library Preparation Kit, 2.5 µL of TD buffer and 1.25 µL of ATM was added to the sample and incubated for 5 minutes at 55 °C. The samples were removed from the heat and had 1.25 µL of NT buffer added, the samples were then incubated at RT for 5 minutes. For the amplification of the libraries and the incorporation of unique indices, 1.25 µL each of an i7 and an i5 index was added as well as 3.75 µL of NPM PCR master mix were added. The libraries were amplified for 12 cycles in a thermocycler using the conditions previously described^67, 68^.

The libraries were purified and size selected using AMPure beads at a 0.85X volume ratio. The AMPure beads were washed twice with 200 µL of 80% ethanol and the libraries were eluted with 20 µL of resuspension buffer. The libraries were sequenced with a NextSeq500 75bp single end reads.

### Preparation of primary myoblast RNA-Seq libraries

MuSCs were FACS sorted as in the “Fluorescence activated cell sorting of muscle stem cells” section described above, and cultured *in vitro* for 3 days in growth media (Ham’s F10, 20% FBS, 1% penicillin/streptomycin, 5 ng/mL bFGF). After 3 days, the cells were trypsinized with 0.125% trypsin and the cells were pelleted at 1500 RPM for 5 minutes. The cells were resuspended in PBS and counted. 1000 cells were transferred to SMART lysis buffer (1 µL SMART-Seq Reaction Buffer (95% SMART-Seq 10X lysis buffer, 5% RNAse Inhibitor), 9 µL ddH_2_0) at a final volume of 12.5 μL. Myoblast RNA-seq libraries were then generated in the same manner as the MuSCs, described above.

### RNA-Seq analysis

To determine the differential gene expression, mRNA expression was quantified with Kallisto v0.46.1 (parameters: --single --fragment-length 200 --sd 20)^69^. The GENCODE v.M28 basic gene annotations was used for transcriptome information^70^. Transcript-level abundance estimates were collapse to the gene level using the “tximport”^71^. Differentially expressed genes were identify using DESeq2^72^. P values were adjusted for multiple testing using the Independent Hypothesis Weighting procedure^73^.

Reads were aligned with HISAT2 and converted to bigwigs using bamCoverage (version 3.5.1). Read counts were normalized as RPM^74, 75^.

### Analysis of differential mRNA stability

Differential mRNA stability was inferred as previously described^76^. Briefly, exonic and intronic regions from the RNA-seq reads were mapped using annotations acquired from Ensembl GRCm39 version 87. The decoupling of the transcriptional and post-transcriptional effects was achieved with DiffRAC determining changes in mRNA stability.

### Extraction and digestion of proteins for total proteomic analyses

Proteins from FACS sorted mouse muscle stem cell populations were extracted in lysis buffer containing 5% sodium dodecyl sulfate (SDS), 100 mM TRIS pH 7.8. Samples were subsequently heated to 99°C for 10 minutes. The lysate was clarified by centrifugation 14,000 x g for 5 minutes. An aliquot corresponding to 10% of the total volume of lysate was diluted to <1% SDS and used for estimation of protein concentration by bicinchoninic acid assay (BCA). In the remaining sample, protein disulfide bonds were reduced by the addition of tris(2-carboxyethyl)phosphine (TCEP) to a final concentration of 20 mM and incubated at 60°C for 30 minutes. Free cysteines were alkylated using iodoacetamide at a final concentration of 30 mM and subsequent incubation at 37°C for 30 minutes in the dark. An equivalent of 10 µg of total protein was used for proteolytic digestion using suspension trapping (STRAP). In brief, proteins were acidified through the addition of phosphoric acid to a final concentration of 1.3% v/v. The sample was subsequently diluted 6-fold in STRAP loading buffer (9:1 methanol: water in 100 mM TRIS, pH 7.8) and loaded onto an S-TRAP Micro cartridge (Protifi LLC, Huntington NY) and spun at 4000 x g for 2 minutes. Samples were washed three times using 150 µL of STRAP loading buffer. Proteins were then proteolytically digested using trypsin at a 1:10 enzyme to substrate ratio for 2 hours at 47°C. Peptides were sequentially eluted in 50 mM ammonium bicarbonate, 0.1% formic acid in water, and 50% acetonitrile. Peptide containing samples were then desalted using self-made R3-STAGE tips. Desalted peptides were vacuum concentrated and reconstituted in 0.1% trifluoroacetic acid (TFA) prior to analysis by LC-MS/MS.

### LC-MS/MS acquisition and data analysis

Peptide containing samples were analyzed by data dependent acquisition (DDA) using an Easy-nLC 1200 online coupled to a Q Exactive Plus. Samples were loaded onto the precolumn (Acclaim PepMap 100 C18, 3 µm particle size, 75 µm inner diameter x 2 cm length) in 0.1% formic acid (buffer A). Peptides were separated using a 100-min binary gradient ranging from 3-40% of buffer B (84% acetonitrile, 0.1% formic acid) on the main column (Acclaim PepMap 100 C18, 2 µm particle size, 75 µm inner diameter x 25 cm length) at a flow rate of 300 nL/min. Full MS scans were acquired from m/z 350-1,500 at a resolution of 70,000, with an automatic gain control (AGC) target of 1 x 10^6^ ions and a maximum injection time of 50 ms. The 15 most intense ions (charge states +2 to +4) were isolated with a window of m/z 1.2, an AGC target of 2 x 10^4^ and a maximum injection time of 64 ms and fragmented using a normalized higher-energy collisional dissociation (HCD) energy of 28. MS/MS were acquired at a resolution of 17,500 and the dynamic exclusion was set to 40 s. DDA MS raw data was processed with Proteome Discoverer 2.5 (Thermo Scientific) and searched using Sequest HT against a mouse UniProt FASTA database. The enzyme specificity was set to trypsin with a maximum of 2 missed cleavages. Carbamidomethylation of cysteine was set as fixed modification and oxidation of methionine as variable modification. The precursor ion mass tolerance was set to 10 ppm, and the product ion mass tolerance was set to 0.02 Da. Percolator was used to assess posterior error probabilities and the data was filtered using a false discovery rate (FDR) <1% on peptide and protein level. The Minora feature detector node of Proteome Discoverer was used for label free quantitation (LFQ) based on precursor areas. LFQ abundances were scaled (normalized) based on the total peptide amount per sample. Only proteins quantified with at least 1 protein unique peptide were retained in the data summary. In order to estimate the relative amount of a particular protein within the sample we calculated and ranked proteins based on their normalized spectral abundance factor (NSAF) values (https://doi.org/10.1021/pr060161n). Hierarchical clustering was conducted using the normalized protein LFQ abundances as inputs for Instant Clue (http://www.instantclue.uni-koeln.de/).

### Preparation of muscle stem cell ATAC-Seq libraries

The ATAC-seq libraries were generated following the OMNI ATAC-seq protocol^77^. 5000 MuSCs from hindlimbs were sorted using FACS into 30 μL of lysis buffer (10 mM Tris-HCl pH 7.5, 10 mM NaCl, 3 mM MgCl_2_, 0.1% Tween-20, 0.1% NP-40, 0.01% Digitonin) and incubated on ice for 5 minutes and then at room temperature for 3 min. Cells were then washed with 100 μL of wash buffer (10 mM Tris-HCl pH 7.5, 10 mM NaCl, 3 mM MgCl_2_, 0.1% Tween-20) and centrifuged at 800 g for 10 min. 10 μL transposition mix (5 μL of Tagment DNA (TD) buffer, 3.2 μL PBS, 0.89 μL Tn5, 0.1% Tween-20, 0.01% Digitonin and 0.75 μL water) was used to resuspend the cell pellet. The samples were incubated for 20 min at 37 °C and were mixed every 5-7 min. DNA purification was then performed with column purification as described (Qiagen, QIAquick PCR Purification Kit Cat: 28104), eluted into a final volume of 20 μL.

Indices were incorporated by adding 30 μL of the PCR reaction mix (10 μL Q5 buffer, 10 μL Q5 enhancer, 1 μL dNTPs, 2.5 μL i7 index, 2.5 μL i5 index, 3.5 μL nuclease free water, 0.5 μL Q5 High Fidelity DNA polymerase). PCR amplification was performed using the previously described conditions for 12 cycles^78, 79^. The libraries were then size selected and purified with Ampure XP beads at a 1:0.85 (v:v) ratio. After the validation of the libraries with a bioanalyzer, samples were sequenced with NovaSeq6000 Sprime, Paired End (PE) 150bp.

### Analysis of ATAC-Seq data

Adapters were removed from raw ATAC-seq reads using Trimmomatic (version 0.39, parameters: LEADING:10 TRAILING:10 MINLEN:50)^80^. Paired reads were aligned to mm10 using BWA-MEM (version 0.7.17)^81^. Samtools was used to index and filter (version 1.16.1, mapping quality > 30)^82, 83^. Duplicated reads were removed using Picard (version 2.26.3)^84^.

ATAC-seq peaks were called with MACS2 (version 2.2.7.1, parameters: --nomodel --shift -100 -- extsize 200 –broad, --gsize mm)^85, 86^.

### Isolation and culture of EDL myofibers

The skin from the hindlimb was removed and the TA muscle was dissected out to provide access to the EDL. The EDL was dissected by cutting the distal and proximal tendon. The EDL was placed in a 1.5 mL tube containing 800 µL of digestion buffer (1000 U/mL collagenase in unsupplemented DMEM) for 1 hour at 37 °C and 5% CO_2_. Using a large bore glass pipette, coated with 10% HS in DMEM, the EDL was transferred to a 6-well plate with 2 mL of unsupplemented DMEM in each well. The wells had previously been coated with 10% HS in DMEM for 30 minutes. The EDL muscle was gently triturated with a coated large bore glass pipette to disassociate the myofibers. The myofibers were then either fixed immediately for the 0 hour time point, or allowed to rest in the unsupplemented DMEM for 1 hour in a cell incubator, set to 37°C and 5% CO_2_, before having the DMEM replaced with growth media (DMEM supplemented with 15% FBS, 1% chick embryo extract, 1% penicillin/streptomycin, 5 ng/mL bFGF).

### Immunofluorescence of EDL myofibers

Isolated EDL myofibers were transferred using a small-bore glass pipette, coated with 10% HS in DMEM, to a 24-well plate. The myofibers were fixed with 400 µL of 4% PFA in PBS for 5 minutes at room temperature. The myofibers were washed three times in 400 µL of 0.1% Triton-X in PBS. Blocking and permeabilization were performed in tandem using 400 µL of blocking buffer (1% Triton-X, 10% goat serum, 6% horse serum, 2% BSA, and 0.1 M of glycine in PBS) for 1 hour at RT. Myofibers were incubated with primary antibodies diluted in blocking buffer overnight at 4 °C. The next day, the samples were washed three times in 400 µL of 0.1% Triton-X in PBS. Next, the samples were incubated with secondary antibody diluted in blocking buffer for 1 hour at RT. The myofibers were then washed three times in 400 µL of 0.1% Triton-X in PBS and afterwards transferred to a glass slide with mounting solution containing DAPI.

### Quantification and statistical analysis

Statistical analyses were performed using Prism9 (Graphpad). Unpaired two-tailed t-tests were used to analyze the difference between experimental conditions. Error bars represent the standard deviation. Sample sizes for each experiment is described in the corresponding figure legend.

## Materials availability

All new reagents and material generated in this study are available upon request.

## Supplementary Information

**Supplemental Figure 1:**
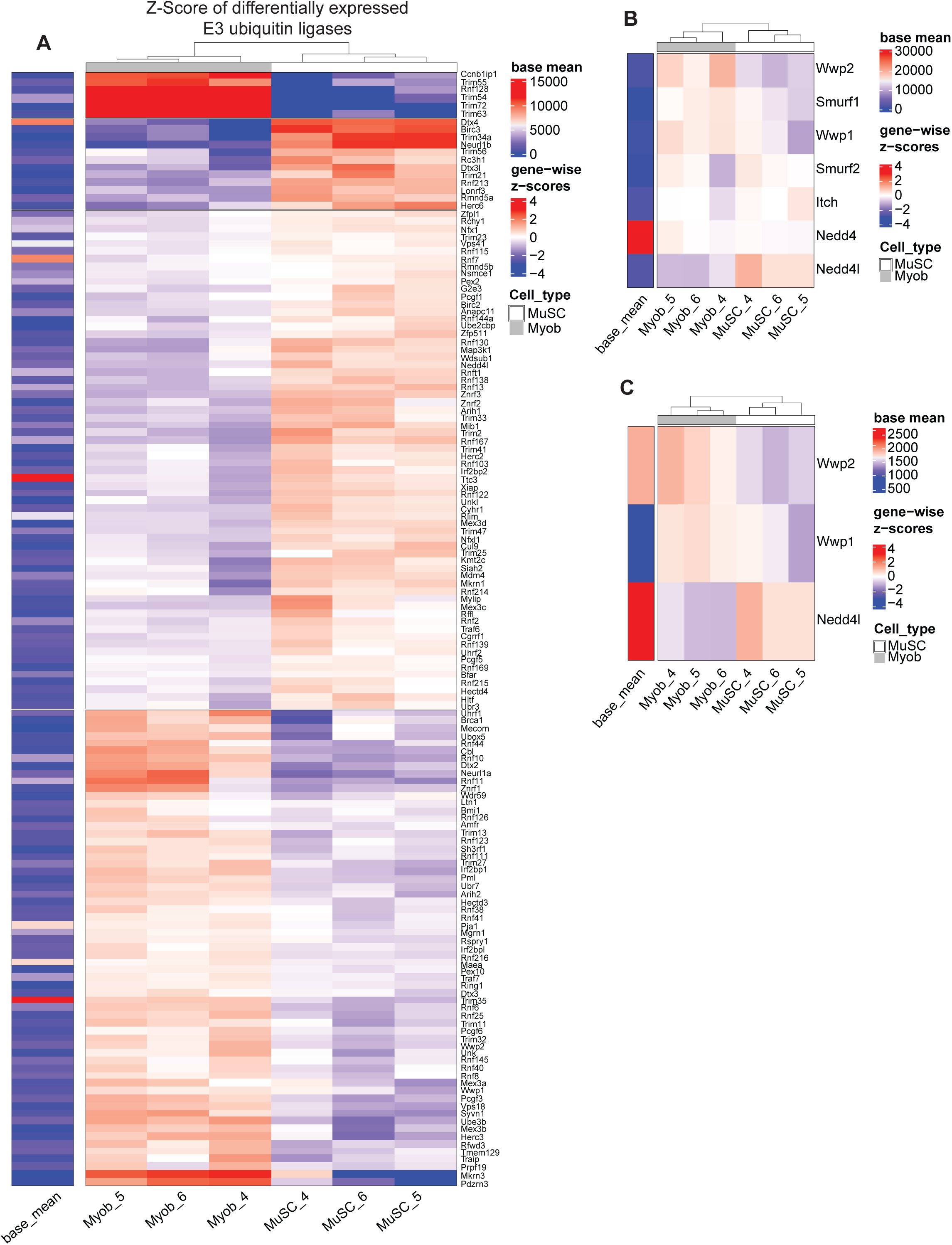
**A.** Heatmap representing the significantly differentially expressed E3 ubiquitin ligases between muscle stem cells and primary myoblasts cultured for 3 days. Genes with a base mean count below 100 were removed. **B.** Heatmap representing the Nedd4 E3 ubiquitin ligase family members expression between muscle stem cells and primary myoblasts cultured for 3 days. **C.** Heatmap representing the significantly differentially expressed Nedd4 E3 ubiquitin ligase family members between muscle stem cells and primary myoblasts cultured for 3 days.

**Supplemental Figure 2:**
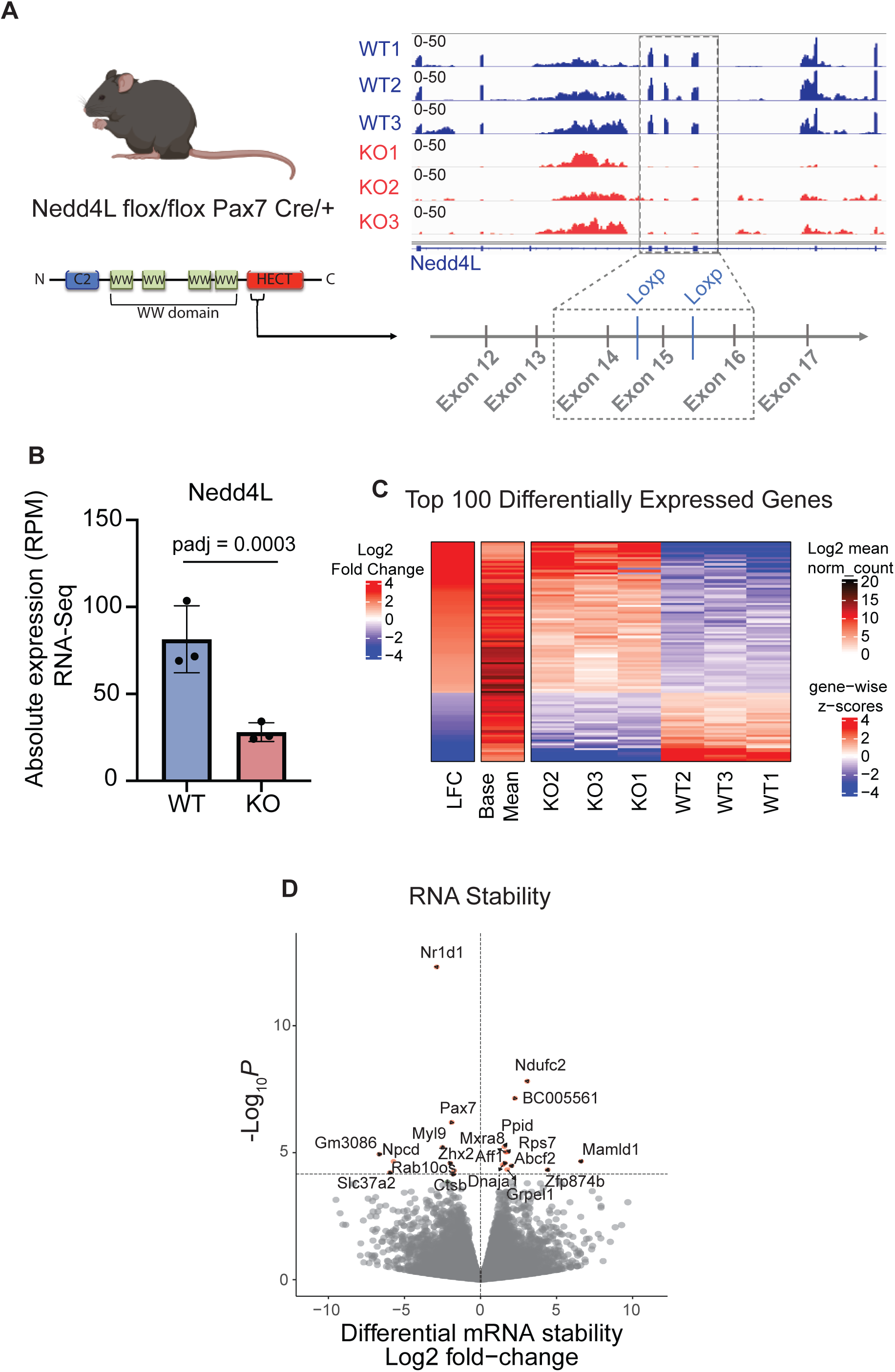
**A.** Diagram representing the genetic mouse model where loxp sites are present, surrounding exon 15, generated with Biorender.com. The cre recombinase is linked to Pax7 expression, resulting in muscle stem cells having exon 15 deleted. **B.** Bar graph representing the absolute expression in RPM from WT and Nedd4L-cKO MuSC RNA-Seq. **C.** Heatmap of the top 100 differentially expressed genes between WT and Nedd4L-cKO MuSCs. **D.** Volcano plot of the RNA stability of transcripts between WT and Nedd4L-cKO MuSCs. Data presented as mean ± SD.

**Supplemental Figure 3:**
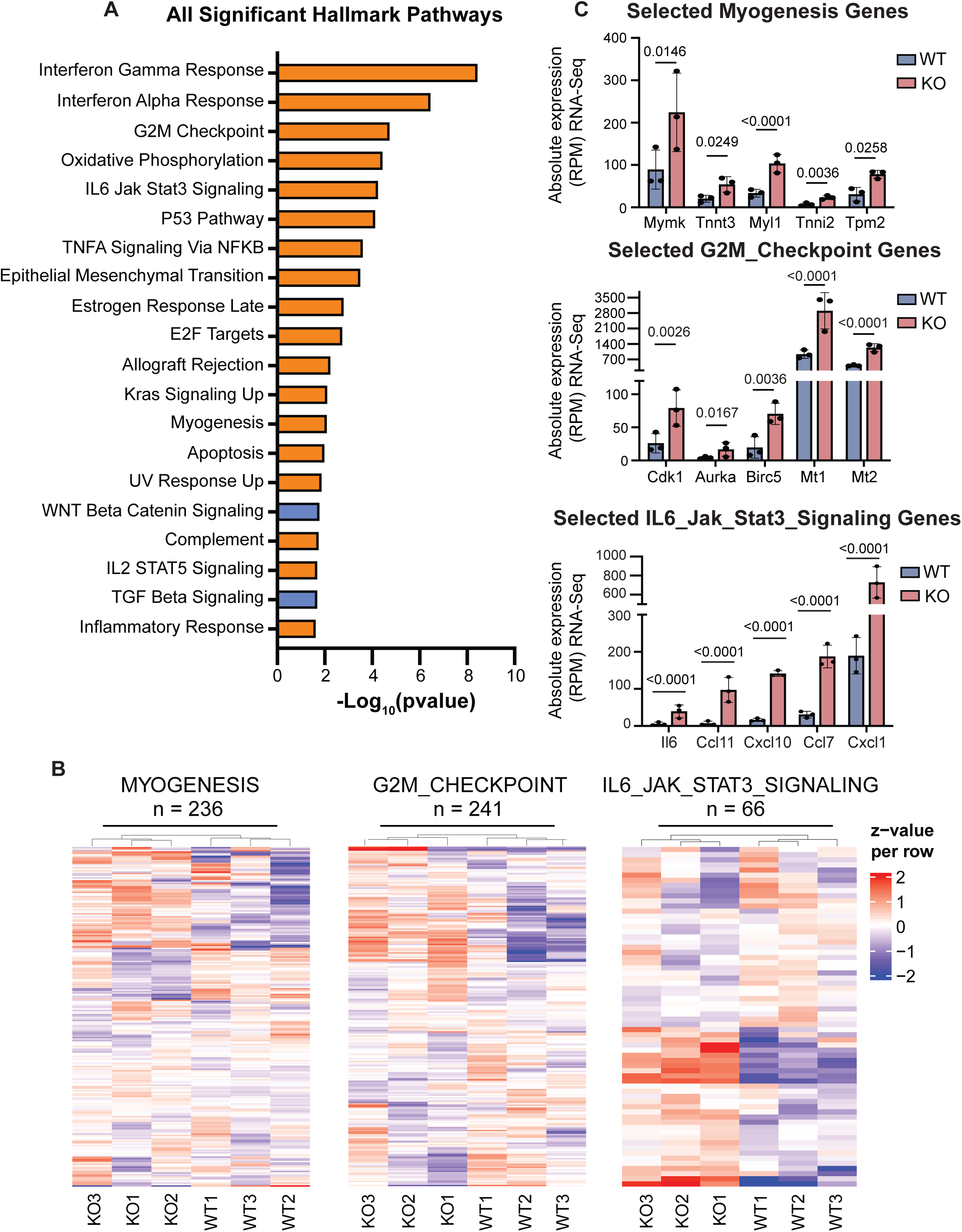
**A.** Bar graph of all of the significantly altered Hallmark pathways between the transcriptome of WT and Nedd4L-cKO MuSCs. Orange bars represent an upregulation in the Nedd4L-cKO; blue bars represent a downregulation. **B.** Heatmaps of the expression of the genes associated with the G2M checkpoint, Myogenesis, and Il6 JAK STAT3 Signaling pathways, in the individual RNASeq replicates. **C.** Bar graphs of selected representative genes in the G2M Checkpoint, Myogenesis, and Il6 JAK STAT3 Signaling pathways. Data presented as mean ± SD.

**Supplemental Figure 4:**
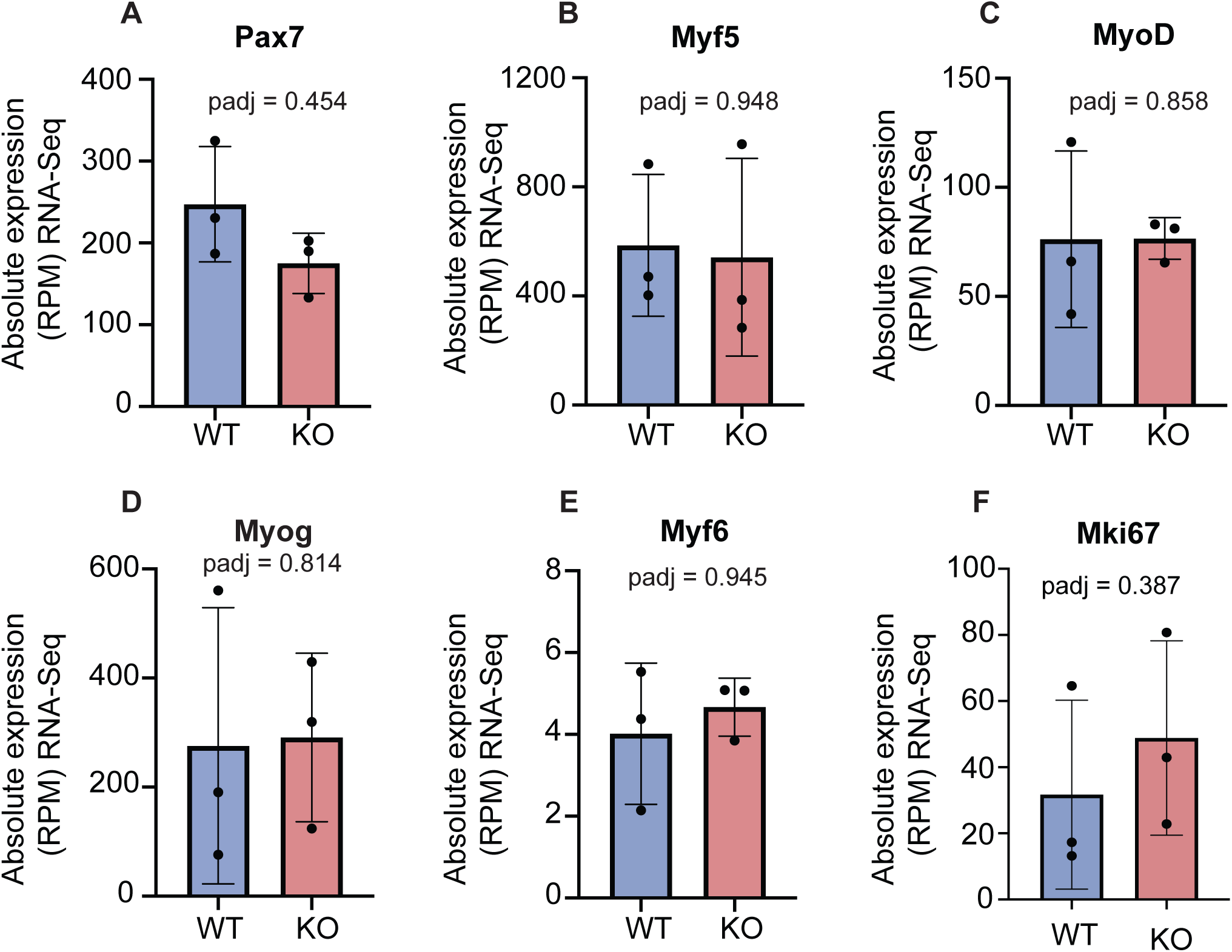
**A.** Bar graph of the absolute expression in RPM of Pax7 from the WT and Nedd4L-cKO MuSC RNASeq. **B.** Bar graph of the absolute expression in RPM of Myf5 from the WT and Nedd4L-cKO MuSC RNA-Seq. **C.** Bar graph of the absolute expression in RPM of MyoD from the WT and Nedd4L-cKO MuSC RNA-Seq. **D.** Bar graph of the absolute expression in RPM of Myogenin from the WT and Nedd4L-cKO MuSC RNA-Seq. **E.** Bar graph of the absolute expression in RPM of Myf6 from the WT and Nedd4L-cKO MuSC RNA-Seq. **F.** Bar graph of the absolute expression in RPM of mki67 from the WT and Nedd4L-cKO MuSC RNA-Seq. Data presented as mean ± SD.

**Supplemental Figure 5:**
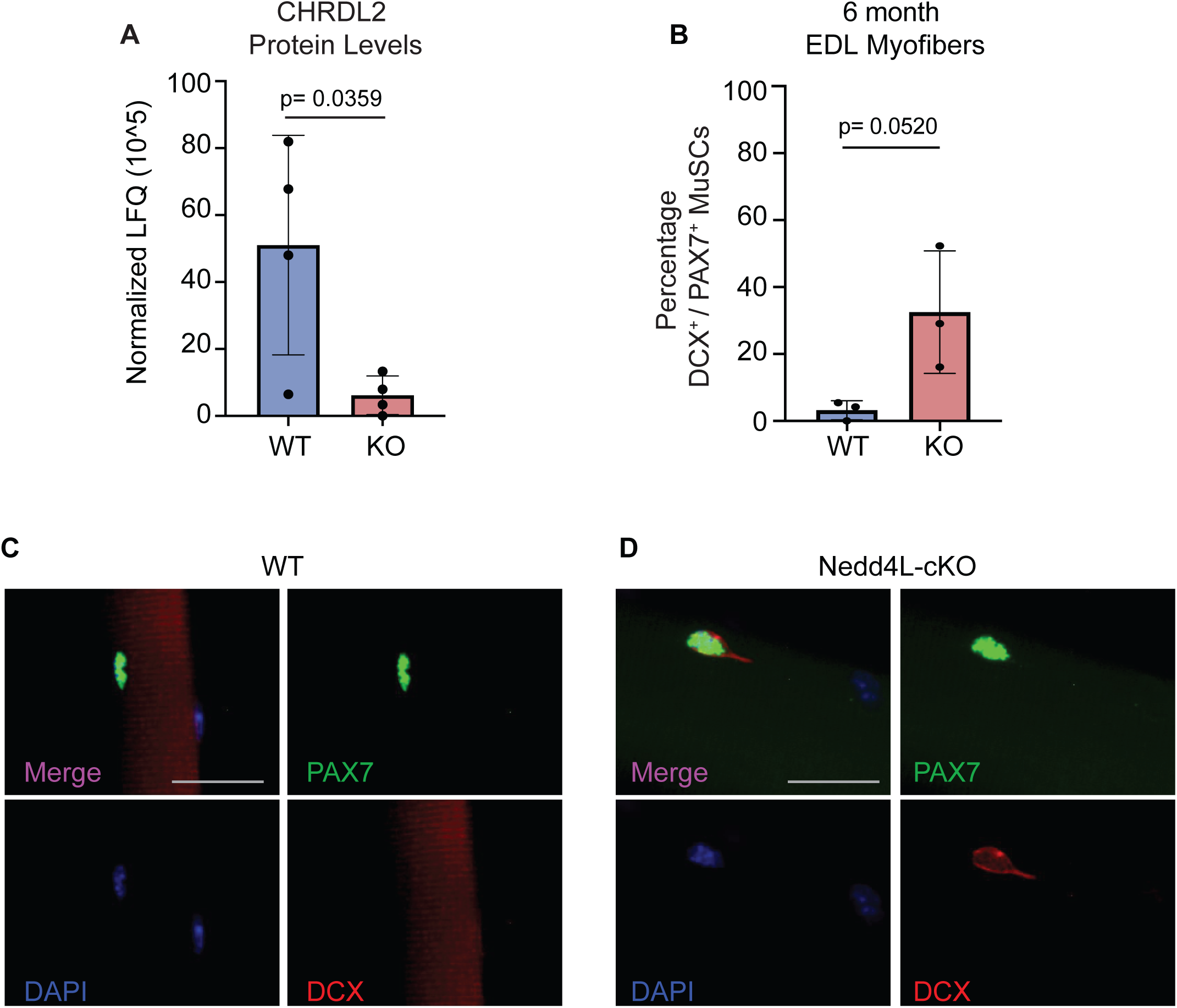
**A.** Bar graph of the CHRDL2 protein levels of WT and Nedd4L-cKO MuSC proteomics. **B.** Bar graph of the percentage of DCX expressing MuSCs associated to 0 hour post isolated EDL myofibers, from 6-month-old male mice. **C.** Representative image of freshly isolated EDL myofibers stained for DCX and PAX7 from WT 6-month-old mice. Scale bar = 25 µm. **D.** Representative image of freshly isolated EDL myofibers stained for DCX and PAX7 from Nedd4L-cKO 6-month-old mice. Scale bar = 25µm. Data presented as mean ± SD.

**Supplemental Figure 6:**
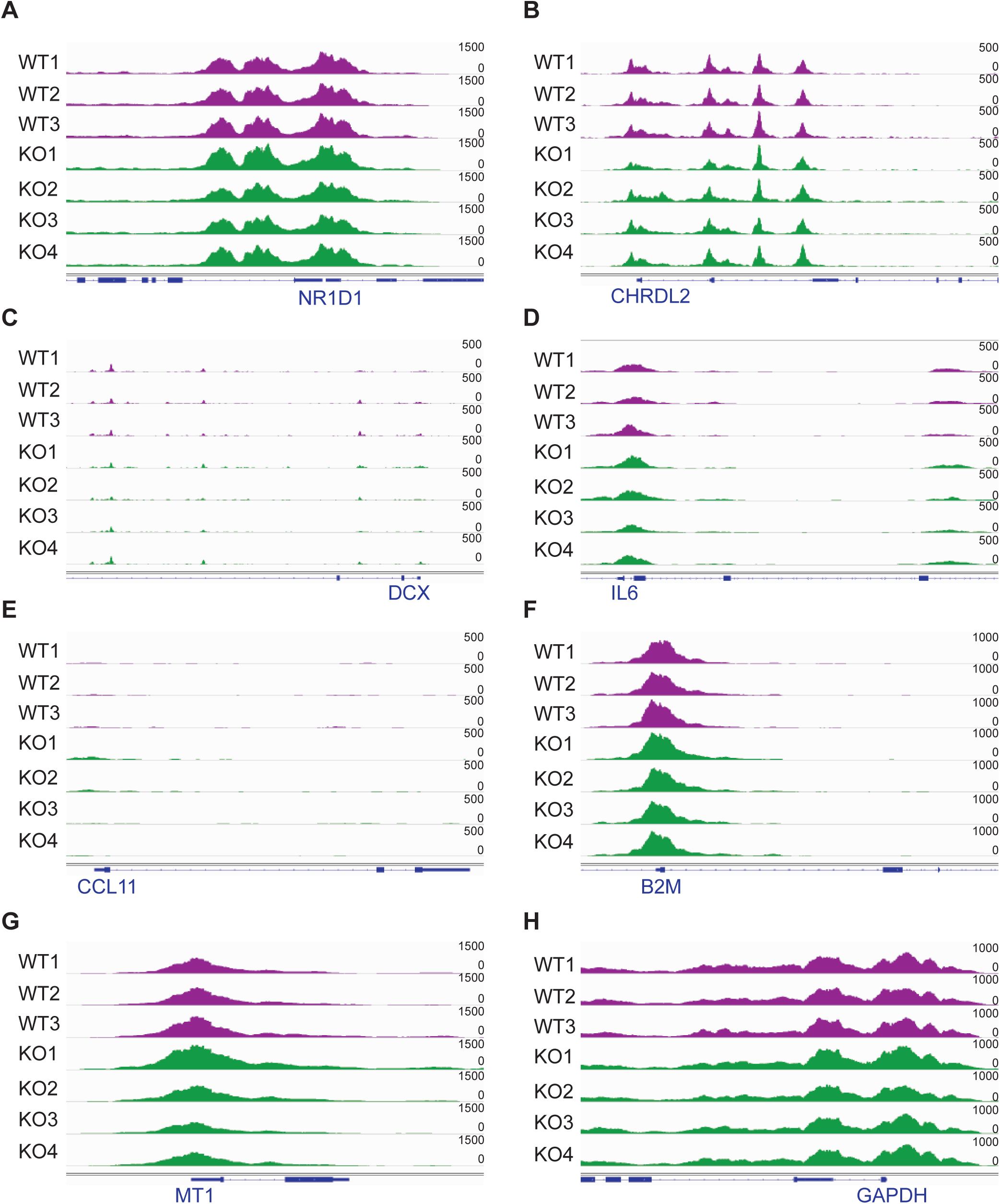
**A.** IGV tracks of ATAC-Seq peaks from WT and Nedd4L-cKO MuSCs of the NR1D1 gene. **B.** IGV tracks of ATAC-Seq peaks from WT and Nedd4L-cKO MuSCs of the CHRDL2 gene. **C.** IGV tracks of ATAC-Seq peaks from WT and Nedd4L-cKO MuSCs of the DCX gene. **D.** IGV tracks of ATAC-Seq peaks from WT and Nedd4L-cKO MuSCs of the IL6 gene. **E.** IGV tracks of ATAC-Seq peaks from WT and Nedd4L-cKO MuSCs of the CCL11 gene. **F.** IGV tracks of ATAC-Seq peaks from WT and Nedd4L-cKO MuSCs of the B2M gene. **G.** IGV tracks of ATAC-Seq peaks from WT and Nedd4LcKO MuSCs of the MT1 gene. **H.** IGV tracks of ATAC-Seq peaks from WT and Nedd4L-cKO MuSCs of the GAPDH gene. Data presented as mean ± SD.

**Supplemental Figure 7:**
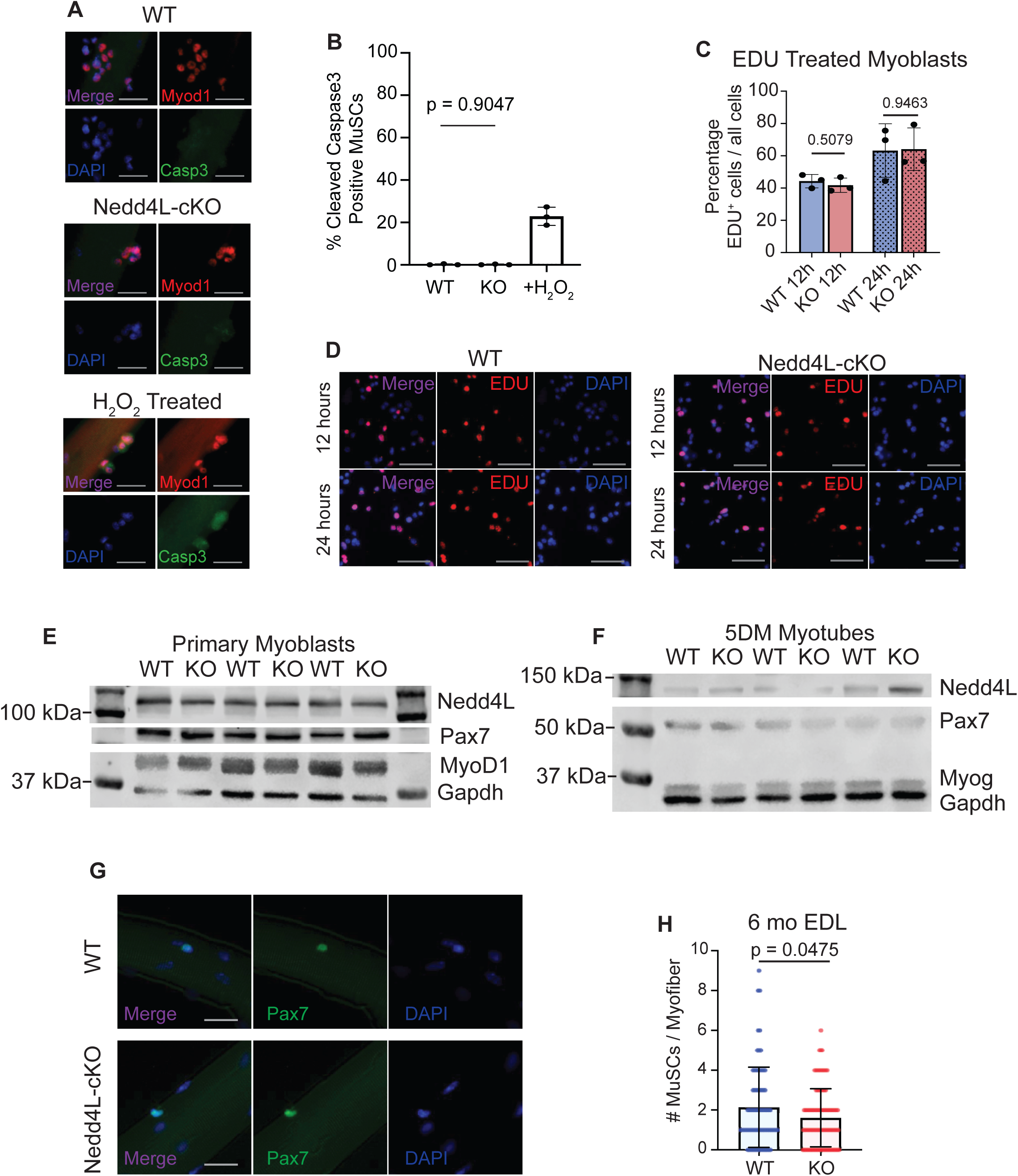
**A.** Representative images of EDL myofibers 72 hours post isolation and stained for MyoD and cleaved Caspase3 from WT, Nedd4L-cKO 6-8 weeks old mice. WT myofibers were treated with H_2_O_2_ as a positive control. Scale bar = 25µm. **B.** Bar graph of the percentage of cleaved Caspase3 positive MuSCs that were associated to EDL myofibers 72 hours post isolation from WT and Nedd4L-cKO myofibers. WT myofibers were treated with H_2_O_2_ as a positive control. **C.** Bar graph of the percentage of EDU positive primary myoblasts treated for 12 and 24 hours. **D.** Representative images of WT and Nedd4L-cKO primary myoblasts treated with EDU for 12 and 24 hours. Scale bar = 25µm. **E.** Western blot of Nedd4L, Pax7, MyoD and Gapdh as a loading control, from WT and Nedd4L-cKO primary myoblasts. **F.** Western blot of Nedd4L, Pax7, Myogenin and Gapdh as a loading control, from WT and Nedd4L-cKO myotubes cultured for 5 days in differentiation media. Data presented as mean ± SD. **G.** Representative images of freshly isolated EDL myofibers from WT and Nedd4L-cKO male 6-month-old mice, stained for Pax7. **H.** Bar graph of the number of MuSCs from 0 hour post isolation EDL myofibers from WT and Nedd4L-cKO 6-month-old mice.

